# Early and late chaperone intervention therapy boosts XBP1s and ADAM10, restores proteostasis, and rescues learning in Alzheimer’s Disease mice

**DOI:** 10.1101/2023.05.23.541973

**Authors:** Jennifer M. Hafycz, Ewa Strus, Nirinjini N. Naidoo

## Abstract

Alzheimer’s disease (AD) is a debilitating neurodegenerative disorder that is pervasive among the aging population. Two distinct phenotypes of AD are deficits in cognition and proteostasis, including chronic activation of the unfolded protein response (UPR) and aberrant Aβ production. It is unknown if restoring proteostasis by reducing chronic and aberrant UPR activation in AD can improve pathology and cognition. Here, we present data using an APP knock-in mouse model of AD and several protein chaperone supplementation paradigms, including a late-stage intervention. We show that supplementing protein chaperones systemically and locally in the hippocampus reduces PERK signaling and increases XBP1s, which is associated with increased ADAM10 and decreased Aβ42. Importantly, chaperone treatment improves cognition which is correlated with increased CREB phosphorylation and BDNF. Together, this data suggests that chaperone treatment restores proteostasis in a mouse model of AD and that this restoration is associated with improved cognition and reduced pathology.

**One-sentence summary:** Chaperone therapy in a mouse model of Alzheimer’s disease improves cognition by reducing chronic UPR activity

## INTRODUCTION

Alzheimer’s disease (AD) is a prevalent neurodegenerative disease and is the leading cause of dementia in older adults (Hebert et al., 1995; Jahn, 2013; Morris, 1996). Despite its widespread nature, the causes of AD pathology are not fully understood, and research to uncover these mechanisms is vital. Indeed, AD is oftentimes difficult to diagnose early on and is typically identified following cognitive impairments, which occur after significant disease progression (Holtzman et al., 2011; Jahn, 2013; Tarawneh and Holtzman, 2012). The pathogenesis of many neurodegenerative diseases includes protein dyshomeostasis and corresponding protein aggregation (Hetz and Saxena, 2017). In AD, these aggregations are commonly seen as tau tangles or amyloid-β aggregations, known as plaques which are made up of Aβ-42 fibrils (Koo et al., 1999; Ross and Poirier, 2004; Thal and Fandrich, 2015). Aβ-42 is generated via amyloidogenic cleavage of amyloid precursor protein (APP) by β-site APP cleaving enzyme 1 (BACE1) (Peron et al., 2018; Yuan et al., 2017). Under healthy conditions, APP is an important synaptic stabilizing protein and is cleaved by an α-secretase, a disintegrin and metalloproteinase 10 (ADAM10), in the non-amyloidogenic pathway (Muller et al., 2012; Peron et al., 2018; Yuan et al., 2017). ADAM10 is reduced in AD, which is thought to contribute to the pathogenesis of the disease (Peron et al., 2018; Sogorb-Esteve et al., 2018; Yuan et al., 2017). While the exact detrimental effects of these protein aggregations are unclear, it is thought that Aβ-42 and these aberrant aggregates alter neuronal signaling and thus contribute to cognitive decline (Medeiros et al., 2011; Morley and Farr, 2014).

The maintenance of protein homeostasis, or proteostasis, is critical for the function and survival of the cell. The protein aggregations seen in AD are indicative of an imbalance in proteostasis and an increase in endoplasmic reticulum (ER) stress (Hetz and Saxena, 2017; Hoozemans et al., 2012). ER stress occurs when proteins misfold and aggregate in the ER lumen (Berridge, 2002; Lee et al., 2002; Ron and Walter, 2007; Szegezdi et al., 2006). ER stress in healthy organisms activates the unfolded protein response (UPR), a protein quality control and signal transduction pathway (Hetz et al., 2020). The UPR restores proteostasis by alleviating the protein folding load on the cell through the activation of three distinct ER stress sensors PERK, IRE1, and ATF6. (Hetz et al., 2020; Kaufman, 2002; Koga et al., 2011; Naidoo et al., 2008). Briefly, PERK activation attenuates global protein translation, apart from a few key targets, including activating transcription factor 4 (ATF4) (Hetz et al., 2020). IRE1 activation leads to increased spliced x-box binding protein 1 mRNA and generation of the transcription factor X-Box Binding Protein 1 (XBP1s), which promotes the synthesis of the chaperone protein, BiP (Lee et al., 2003), while ATF6 activation also promotes chaperone synthesis (Hetz et al., 2020). Collectively, these pathways work to resolve ER stress and restore proteostasis.

Importantly, age is a major risk factor for the development of AD, and there is a plethora of evidence that demonstrates increased UPR activity in AD brains, both in post-mortem human tissue as well as in several animal models of AD and other neurodegenerative diseases (Gavilan et al., 2009; Halliday and Mallucci, 2014; Hetz and Saxena, 2017; Hoozemans et al., 2009; Hoozemans et al., 2012). Under healthy aging conditions, the mechanisms that maintain proteostasis are less efficient and can even become maladaptive (Brown et al., 2014; Hafycz et al., 2022; Naidoo et al., 2008; Naidoo et al., 2011). Specifically with age, there is increased ER stress and the UPR is more chronically activated (Brown et al., 2014; Brown and Naidoo, 2012; Hafycz et al., 2022; Hetz and Mollereau, 2014; Koga et al., 2011; Naidoo et al., 2008). Chronic UPR activity promotes pro-apoptotic signaling and cell death via prolonged PERK signaling and subsequent CHOP activation (Hetz and Mollereau, 2014; Koga et al., 2011). With age, there is also a reduction in the endogenous ER protein chaperone BiP, which is crucial to prevent protein misfolding (Brown et al., 2014; Hetz et al., 2020; Naidoo et al., 2008; Naidoo et al., 2011; Paz Gavilan et al., 2006). Further, chronically inhibited protein translation via PERK activation has many consequences, including memory deficits (Hernandez and Abel, 2008; Hughes and Mallucci, 2019; Hussain and Ramaiah, 2007; Sen et al., 2017; Sharma et al., 2018). Interestingly, the UPR can modulate ADAM10 levels, as XBP1s, downstream of IRE1, acts as a transcription factor to promote ADAM10 (Cisse et al., 2017; Reinhardt et al., 2014). However, it remains unclear if UPR activation is a symptom of AD or if chronic maladaptive UPR activation is causal to the progression of the disease. Further, it is unknown if modulating the UPR can alter XBP1s levels, and thus ADAM10, to improve AD pathology.

We have shown previously, in both a *drosophila* model and a mouse model of aging, that supplementing chaperone levels with a small chemical chaperone, 4-phenylbutyrate (PBA) reduces chronic UPR activity in the brain (Brown et al., 2014; Hafycz et al., 2022). Here, we wanted to determine if UPR activity plays a similar role in cognition and pathology in an amyloid precursor protein (APP) knock in mouse model of AD. We used the *APP^NL-G-F^* model, developed to include the Swedish, Iberian, and Arctic mutations, and which uses the endogenous mouse APP promoter to drive the expression of APP (Saito et al., 2014). We employed three distinct groups of these *APP^NL-G-F^* knock-in (KI) mice and their *APP^WT^* wildtype (WT) littermates to determine if intervening at an early age or at a later age, when the disease has progressed, can improve behavior and pathology.

We report that chaperone treatment improved cognitive performance with both early and late-stage intervention in the *APP^NL-G-F^* KI mice. Chaperone treatment also led to improved proteostasis, with a diminution of ER stress in the hippocampus and an increase in ADAM10 and CREB activation in *APP^NL-G-F^*KI mice. Promisingly, hippocampal BiP overexpression recapitulated these results. Altogether, our data demonstrate that disrupted proteostasis and chronic UPR activity play a key role in AD, and that supplementing chaperone levels is sufficient to restore both proteostasis and cognition during early and late-stage intervention paradigms. Together, this work could inform the development of novel therapeutics to help treat this debilitating disease.

## RESULTS

### 1: Early PBA treatment improves proteostasis in the hippocampus of *APP^NL-G-F^* KI mice

Given the prevalence of UPR activity in brain tissues from AD patients and animal models (Hetz and Saxena, 2017; Hoozemans et al., 2009; Hoozemans et al., 2012), and the fact that aberrant UPR activity, present in aging, is coupled with diminished BiP levels (Brown et al., 2014; Hafycz et al., 2022; Naidoo et al., 2008), we first wanted to establish if BiP is inherently reduced in *APP^NL-G-F^* KI mice. Since the *APP^NL-G-F^*mutations are constitutive, we intervened with intraperitoneal (I.P.) PBA or saline injections starting from weaning. Following 10 weeks of treatment and behavioral testing, we used immunofluorescence to probe for BiP and subsequent markers of UPR activity.

#### 1.1 Early PBA treatment restores BiP levels in the APP^NL-G-F^ KI mice

We found that BiP levels are reduced in the hippocampus of saline treated *APP^NL-G-F^* KI mice, most notably in the CA3 region (p<0.001, Fig. 1A), but also in the CA1 and dentate gyrus (p<0.01 and p<0.05, respectively; Supplemental Fig. 1A-B, Supplemental Table 1) compared to saline treated WT littermate controls. Following PBA treatment, the *APP^NL-G-F^* KI mice displayed restored levels of BiP in the CA3 (p<0.01; Fig. 1A, Supplemental Table 1), as well as in the dentate gyrus (p=0.05; Supplemental Fig. 1A-B, Supplemental Table 1), indicative of an increase in endogenous chaperone synthesis with PBA treatment.

**Fig. 1:**
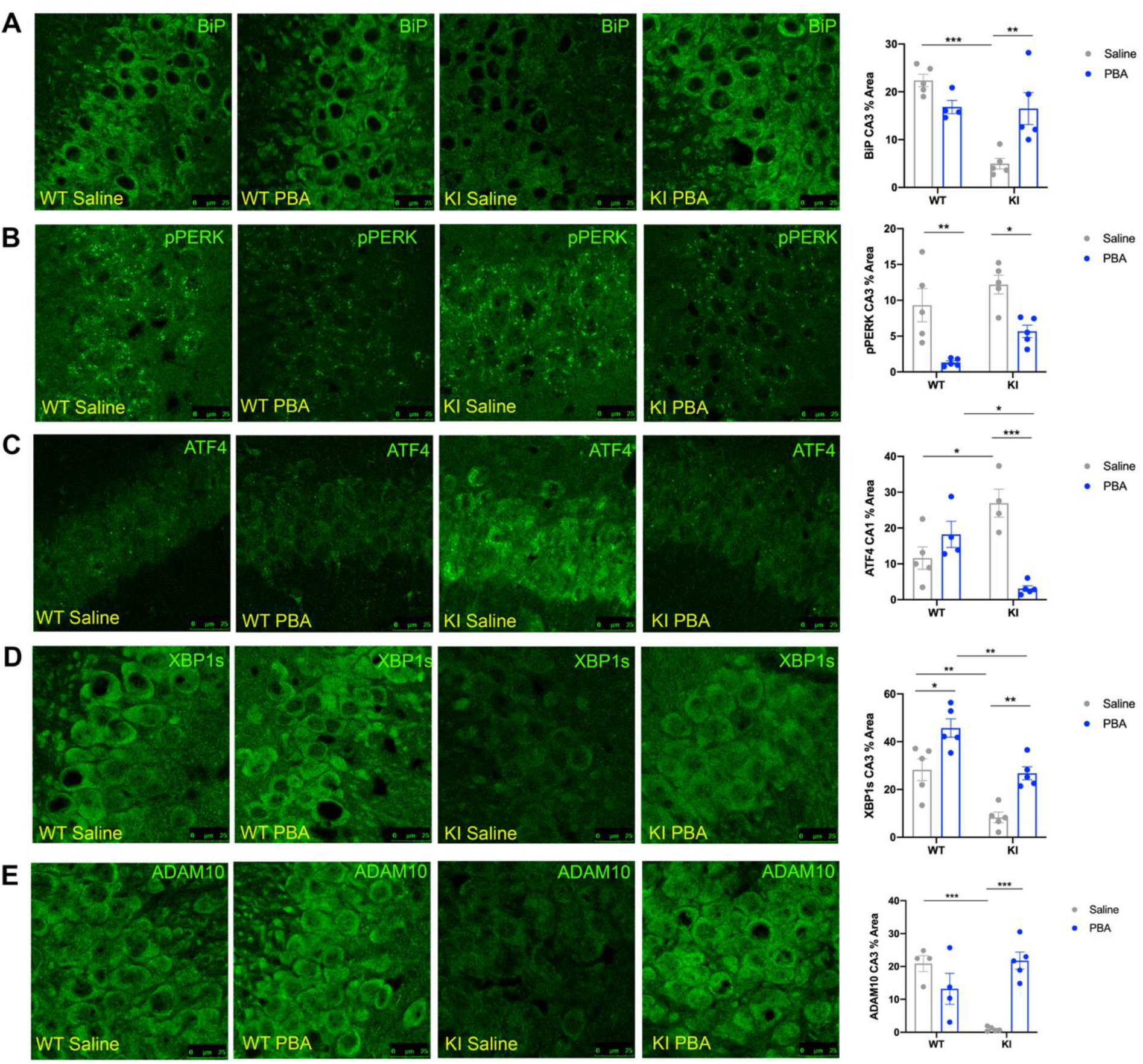
PBA treatment reduces PERK activity and increases XBP1s in the hippocampus of *APP^NL-G-F^* KI mice. Confocal images of the hippocampus across groups. **A**: BiP in the CA3; **B**: p-PERK in the CA3; **C**: ATF4 in the CA1; **D**: XBP1s in the CA3; **E**: ADAM10 in the CA3. Data quantified and presented as mean ± SE percent area of BiP, p-PERK, ATF4, XBP1s, and ADAM10 within hippocampal sections (n=4-5 animals per group; two-way ANOVA with Tukey post-hoc correction for multiple comparisons, *p<0.05, **p<0.01, ***p<0.001).

#### 1.2 Early PBA treatment reduces PERK signaling

Using immunofluorescence to assess PERK activation, we found no difference in p-PERK levels in the CA3 region of the hippocampus in *APP^NL-G-F^* KI mice when compared to *APP^WT^* WT mice under saline-treated conditions (p>0.4; Fig. 1B, Supplemental Table 1). Both *APP^NL-G-F^* KI and *APP^WT^* WT PBA-treated mice showed a decrease in PERK phosphorylation in the CA3 compared to saline treated mice (p<0.05 and p<0.01, respectively; Fig. 1B, Supplemental Table 1), demonstrating that PBA treatment can reduce baseline levels of UPR activity independent of genotype.

Next, we assessed levels of activating transcription factor 4 (ATF4), which is a transcription factor that bypasses the translation block induced by p-PERK phosphorylation of eIF2α. *APP^NL-G-F^* KI mice displayed more ATF4 staining in the CA1 than *APP^WT^* WT littermates (p<0.05 Fig. 1C, Supplemental Table 1). *APP^NL-G-F^* KI PBA-treated mice had less ATF4 in the CA1 relative to saline treated *APP^NL-G-F^* KI mice (p<0.001; Fig. 1C, Supplemental Table 1). This suggests that PBA treatment from weaning reduces ATF4, which is downstream of the PERK-peIF2α translation block, in the hippocampi of *APP^NL-G-F^* KI mice.

#### 1.3 Early PBA treatment in APP^NL-G-F^ KI mice increases XBP1s and ADAM10

In addition to PERK signaling, we also examined activation of the IRE1, which leads to XBP1s synthesis. XBP1s contributes to an adaptive pro-survival UPR response, as XBP1s promotes chaperone synthesis (Hetz et al., 2020; Lee et al., 2003). We hypothesized that *APP^NL-G-F^* KI mice exhibit more chronic, maladaptive UPR activity and thus display a diminished adaptive response. Using immunofluorescence, we found that saline-treated *APP^NL-G-F^* KI mice had less XBP1s relative to WT littermates in the CA3 of the hippocampus (p<0.01; Fig. 1D, Supplemental Table 1), as well as in the dentate gyrus (p<0.05; Supplemental Fig. 1D). PBA treatment from weaning increased XBP1s in both *APP^NL-G-F^* KI and *APP^WT^*WT mice in the CA3 (p<0.01, p<0.05 respectively; Fig. 1D, Supplemental Table 1), as well as in the CA1 of *APP^NL-G-F^* KI and *APP^WT^* WT (p<0.01 and p<0.001, respectively; Supplemental Fig. 1C) and the dentate gyrus (p<0.05 and p<0.05, respectively; Supplemental Fig. 1D), altogether demonstrating that an early intervention with PBA treatment increases an adaptive UPR response via promoting XBP1s.

Interestingly, XBP1s regulates ADAM10, which is an α-secretase that promotes non-amyloidogenic cleavage of APP (Peron et al., 2018; Reinhardt et al., 2014). We probed for ADAM10 to determine if the changes observed in UPR activity are linked to APP processing. We observed less ADAM10 in the CA3 region of the hippocampus in saline-treated *APP^NL-G-F^*KI mice compared to saline treated *APP^WT^* WT mice (p<0.001, Fig. 1E, Supplemental Table 1). With PBA treatment, the *APP^NL-G-F^* KI mice displayed increased ADAM10 levels compared to saline treated *APP^NL-G-F^*KI mice (p<0.001, Fig. 1E, Supplemental Table 1). This data indicates that chaperone treatment could affect how APP is processed via altering the amount of ADAM10, downstream of XBP1s.

### 2: PBA treatment at an early age improves learning in *APP^NL-G-F^* KI mice

Memory deficits are a central phenotype of AD (Alzheimer’s, 2016; Holtzman et al., 2011; Jahn, 2013). Many studies have shown previously that memory deficits are linked to increased ER stress (Hafycz et al., 2022; Moreno et al., 2012; Ricobaraza et al., 2012). To probe cognitive deficits in the *APP^NL-G-F^* KI mice and to establish the effect of PBA treatment on cognition, both male and female *APP^NL-G-F^*KI and *APP^WT^*WT mice were subjected to the Spatial Object Recognition (SOR) cognitive task and the Y-maze task (Fig. 2A). Learning was quantified in the SOR test by calculating a discrimination index between the moved and unmoved objects (Bevins and Besheer, 2006; Cavoy and Delacour, 1993; Hafycz et al., 2022; Sivakumaran et al., 2018), while spatial working memory in the Y-maze test was measured by calculating percent alternations (Hafycz et al., 2022; Kraeuter et al., 2019).

**Fig. 2:**
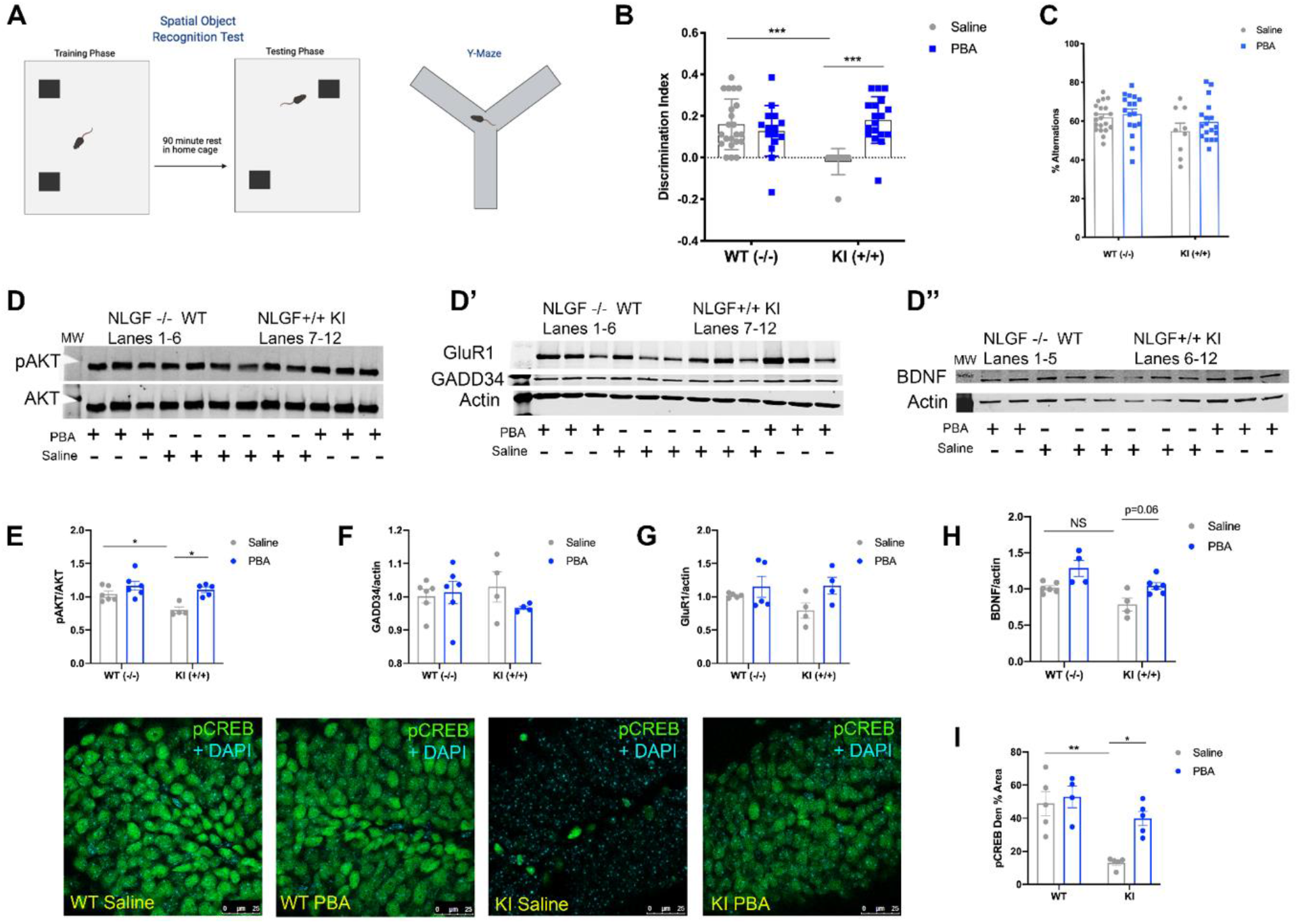
PBA treatment improves learning in the SOR test in *APP^NL-G-F^* KI mice, which is correlated with an increase in the phosphorylation of CREB and BDNF. **A:** Schematic of SOR and Y-maze tests; **B**: Discrimination index of SOR test; **C:** Percent alternations from Y-maze test; **D-D’’:** Representative western blot images; each lane is a different animal. Membranes are cut along the MW markers following blocking to utilize the two-channel Odyssey system and varying molecular weights for the proteins of interest to probe for multiple markers within one membrane. **D:** pAKT and AKT, ∼65 kDa; **D’:** GluR1 ∼110 kDa, GADD34 ∼100 kDa, Actin ∼42 kDa; **D’’:** BDNF dimer ∼28 kDa, Actin ∼42 kDa. **E:** Quantification of western blot analysis for pAKT/AKT; **F:** Quantification of western blot analysis for GADD34/actin; **G:** Quantification of western blot analysis for GluR1/actin; **H:** Quantification of western blot analysis for BDNF/actin. **Bottom**: Confocal images of the hippocampal dentate gyrus across groups; **I**: Mean ± SE percent area of pCREB in the dentate gyrus; n=4-6 animals per group for Western blot and immunohistochemical assays. (All data presented as mean ± SE; all quantifications analyzed via two-way ANOVA with Tukey post-hoc correction for multiple comparisons, *p<0.05, **p<0.01, ***p<0.001).

#### 2.1 Early PBA treatment improves SOR test performance in APP^NL-G-F^ KI mice

We found that *APP^WT^* WT mice were able to discriminate between the moved and unmoved objects in the SOR test regardless of treatment. *APP^NL-G-F^* KI mice performed poorly in this test compared to saline-treated WT littermates, with a clear inability to discriminate between the two objects (p<0.001; Fig. 2B, Supplemental Table 4). With PBA treatment, *APP^NL-G-F^*KI mice showed improved performance, able to distinguish between the moved and unmoved objects (p<0.001; Fig. 2B, Supplemental Table 4). Male and female mice performed similarly within genotype in response to saline and PBA treatment (Supplemental Fig. 4A-B). However, there were no differences observed in performance in the Y-maze test across treatment or genotype (Fig. 2C). Together, this data suggests that PBA treatment improves spatial learning in the *APP^NL-G-F^* KI mice.

#### 2.2 Early PBA treatment increases phosphorylation of CREB and AKT in the hippocampus of APP^NL-G-F^ KI mice

We have shown that chaperone treatment correlates with increased CREB phosphorylation in the dentate gyrus of the hippocampus in aged mice (Hafycz et al., 2022). To determine if a similar mechanism is at work in the *APP^NL-G-F^* KI model, we examined CREB activation. Male and female mice were pooled together for all immunofluorescence assays as SOR performance was not different between males and females. An n=4-5 mice per group, were utilized for immunofluorescence assays to best correlate changes in SOR performance with changes at the cellular level.

Saline treated *APP^NL-G-F^*KI mice displayed reduced levels of p-CREB in the dentate gyrus compared to saline-treated *APP^WT^* WT mice (p<0.01; Fig. 2, Supplemental Table 1). Following PBA treatment *APP^NL-G-F^* KI mice showed an increase in phosphorylated CREB relative to saline-treated *APP^NL-G-F^* KI mice (p<0.05; Fig. 2, Supplemental Table 1).

Previous work has shown that the activation of the CREB kinase, AKT, can be altered by the UPR through GADD34, which is downstream of p-PERK and ATF4 (Farook et al., 2013; Hafycz et al., 2022; Sen, 2019). We next examined *APP^NL-G-F^* KI and WT tissue for p-AKT and GADD34 via western blot assays. We observed less p-AKT in the hippocampus of saline-treated *APP^NL-G-F^* KI mice compared to saline-treated *APP^WT^* WT mice (p<0.05; Fig. 2D, 2E). Following PBA treatment, *APP^NL-G-F^* KI mice displayed increased levels of p-AKT compared to saline treated *APP^NL-G-F^* KI mice (p<0.05; Fig. 2D, 2E). However, we observed no changes in GADD34 across the groups (Fig. 2D’, 2F).

CREB activation is regulated by synaptic activity, specifically, AMPA signaling is known to lead to CREB activation and AMPA levels are known to be altered in AD (Babaei, 2021; Barco et al., 2002; Middei et al., 2013; Qu et al., 2021). Thus, we quantified the AMPA subunit GluR1 via western blot assays in the hippocampus of saline and PBA treated *APP^NL-G-F^*KI and WT mice. We observed no changes in GluR1 levels in any groups, regardless of genotype or treatment (Fig. 2D’, 2G). To further establish if changes in p-CREB levels are related to changes in cognition, we used Western blot assays to probe for BDNF, a growth factor downstream of CREB well-known for its role in memory formation (Bekinschtein et al., 2008; Gonzalez et al., 2019; Panja and Bramham, 2014). Western analyses did not reveal an obvious deficit in BDNF in the *APP^NL-G-F^*KI saline-treated mice relative to *APP^WT^* WT saline-treated mice (Fig. 2D’’, 2H). While PBA treatment did indicate a trend towards increased BDNF levels in the *APP^NL-G-F^* KI mice compared to saline treated *APP^NL-G-F^* KI mice, it was not significant (p=0.06; Fig. 2D’’, 2H). Together, this data suggests that chaperone treatment could improve cognition by increasing p-CREB signaling, and potentially BDNF, in the *APP^NL-G-F^*KI mice via modulating p-AKT activation.

### 3: Late-stage PBA intervention reduces maladaptive UPR activity and improves learning in older *APP^NL-G-F^* KI mice

As AD is generally not diagnosed until after serious cognitive deficits have manifested (Holtzman et al., 2011; Tarawneh and Holtzman, 2012), we carried out a late-stage PBA intervention paradigm on a second group of *APP^NL-G-F^* KI and *APP^WT^* WT mice starting at 10-12 months of age and continuing for 10-12 weeks.

#### 3.1 Late-stage PBA intervention increased BiP and reduced CHOP levels

Probing for BiP, we found that these aged saline-treated *APP^NL-G-F^* KI mice also exhibited reduced levels of BiP in the CA3 and dentate gyrus of the hippocampus compared to saline treated *APP^WT^* WT mice (p<0.001 and p<0.05, respectively; Fig. 3A, Supplemental Fig. 3A, Supplemental Table 3). BiP levels were restored with PBA treatment in the *APP^NL-G-F^* KI mice in the CA3 and dentate (p<0.01 and p<0.05, respectively; Fig. 4, Supplemental Fig. 3A, Supplemental Table 3). PERK phosphorylation and ATF4 levels were not altered between the aged *APP^NL-G-F^*KI and *APP^WT^* WT mice regardless of treatment (immunofluorescence images not shown; Supplemental Table 3), suggesting that delaying chaperone treatment is not sufficient to combat exacerbated PERK phosphorylation. We also probed for CHOP in the hippocampus of these late-stage PBA intervention mice. CHOP is a transcriptional target of ATF4 and is known to induce pro-apoptotic signaling (Hetz et al., 2020; Kaufman, 2002; Rozpedek et al., 2016). We found that saline treated *APP^NL-G-F^*KI mice had more CHOP in the CA3 region of the hippocampus than saline treated *APP^WT^* WT mice (p<0.001; Fig. 3B, Supplemental Table 3). PBA-treated *APP^NL-G-F^*KI mice displayed reduced CHOP in the CA3 relative to saline-treated *APP^NL-G-F^* KI mice (p<0.01; Fig. 3B, Supplemental Table 3). This indicates that while upstream PERK activity is not altered with PBA treatment, downstream pro-apoptotic signaling via the PERK-ATF4-CHOP pathway is reduced with PBA in *APP^NL-G-F^*KI mice.

**Fig. 3:**
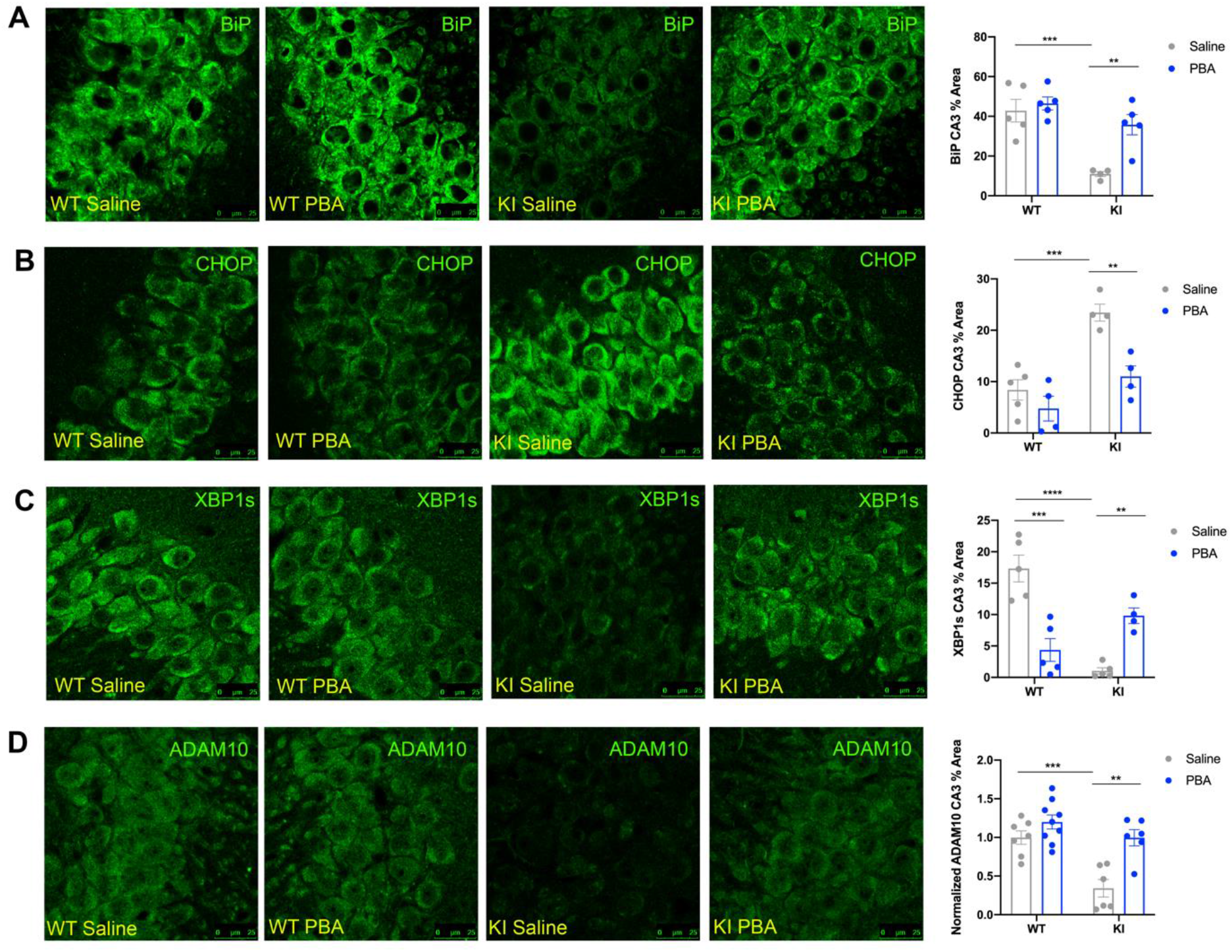
Late-stage PBA treatment in 10–12-month-old *APP^NL-G-F^* KI mice improves adaptive UPR activity in the hippocampus. Representative confocal images of the hippocampus across groups. **A**: BiP immunofluorescent images in the CA3; **B**: CHOP immunofluorescent images in the CA3; **C**: XBP1s immunofluorescent images in the CA3; **D**: ADAM10 immunofluorescent images in the CA3. (Data presented as mean ± SE percent area of BiP, CHOP, XBP1s, and ADAM10 immunofluorescence within hippocampal sections; n=4-9 animals per group; two-way ANOVA with Tukey post-hoc correction for multiple comparisons, *p<0.05, **p<0.01, ***p<0.001, ****p<0.0001).

**Fig. 4:**
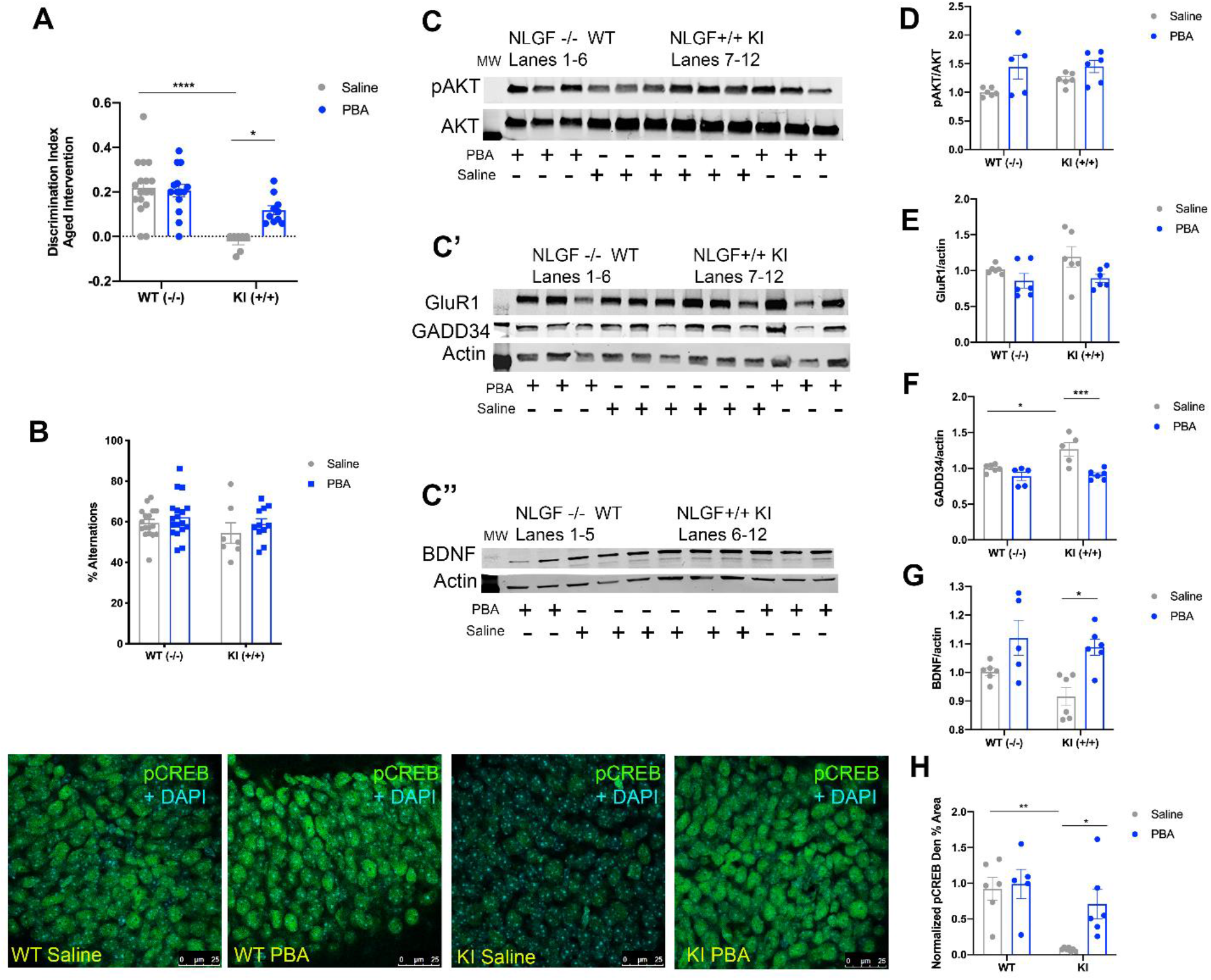
Late-stage PBA intervention improves learning in the SOR test in *APP^NL-G-F^* KI mice, reduces GADD34, and increases CREB and BDNF levels. **A:** Discrimination index of SOR test; **B:** Percent alternations in the Y-maze test; **C-C’’:** Representative western blot images; each lane is a different animal. Membranes are cut along the MW markers following blocking to utilize the two-channel Odyssey system and varying molecular weights for the proteins of interest to probe for multiple markers within one membrane. **C:** pAKT and AKT, ∼65 kDa; **C’:** GluR1 ∼110 kDa, GADD34 ∼100 kDa, Actin ∼42 kDa; **C’’:** BDNF dimer ∼28 kDa, Actin ∼42 kDa. **D:** Quantification of western blot analysis for pAKT/AKT; **E:** Quantification of western blot analysis for GADD34/actin; **F:** Quantification of western blot analysis for GluR1/actin; **G:** Quantification of western blot analysis for BDNF/actin. **Bottom Right**: Confocal images of the hippocampal dentate gyrus across groups; **H**: Quantification of pCREB immunofluorescence normalized to *APP^WT^*WT mice, mean ± SE percent area within hippocampal dentate gyrus sections. n=5-6 animals per group for Western blot and immunohistochemical assays. (All data presented as mean ± SE; all quantifications analyzed via two-way ANOVA with Tukey post-hoc correction for multiple comparisons, *p<0.05, **p<0.01, ***p<0.001).

#### 3.2 Late-stage PBA treatment increased XBP1s and ADAM10 in APP^NL-G-F^ KI mice

In addition, we examined XBP1s levels and found that saline-treated *APP^NL-G-F^* KI mice displayed less XBP1s in the CA3 of the hippocampus relative to saline-treated *APP^WT^* WT mice (p<0.0001; Fig. 3C, Supplemental Table 3), as well as in the CA1 (p<0.01; Supplemental Fig. 2). With PBA treatment, *APP^NL-G-F^* KI mice exhibited restored levels of XBP1s in the CA3 and CA1 compared to saline treated *APP^NL-G-F^* KI mice (p<0.01 and p<0.01, respectively; Fig. 3C, Supplemental Fig. 2, Supplemental Table 3). Together, this data demonstrates that even a late-stage chaperone intervention can reduce chronic UPR signaling and restore adaptive UPR functions.

To determine if APP processing could be altered with PBA treatment, we probed for ADAM10 in these late-stage intervention mice. We found that there was less ADAM10 staining in the CA3 of saline-treated *APP^NL-G-F^* KI mice compared to *APP^WT^* WT mice (p<0.001; Fig. 3D, Supplemental Table 3). With PBA treatment, ADAM10 was increased in the CA3 of *APP^NL-G-F^* KI mice relative to saline treated *APP^NL-G-F^*KI mice (p<0.01; Fig. 3D, Supplemental Table 3). This shows that a late-stage chaperone intervention leads to increased ADAM10 through modulation of the UPR response.

#### 3.3 PBA intervention at a later stage improved learning in the APP^NL-G-F^ KI mice

To determine if late-stage chronic PBA treatment can improve learning in the *APP^NL-G-F^* KI mice, following 10 weeks of either saline or PBA injections, all mice were subjected to the SOR test. As before, saline treated *APP^NL-G-F^* KI mice were unable to distinguish between the moved and unmoved objects, compared to saline treated *APP^WT^*WT mice (p<0.0001; Fig. 4A, Supplemental Table 4). With PBA treatment, the *APP^NL-G-F^* KI mice were able to distinguish between the objects, indicative of improved learning (p<0.05; Fig. 4A, Supplemental Table 4). We observed no differences between males and females in this learning test (Supplemental Figure 4E-F). We also observed no differences in percent alternations in the Y-maze test across groups (Fig. 4B). Promisingly, overall, this indicates that a late-stage chaperone treatment intervention can improve learning in the *APP^NL-G-F^* KI mice.

#### 3.4 Late-stage PBA intervention increased CREB phosphorylation and BDNF levels in the APP^NL-G-F^ KI mice

Next, we measured CREB phosphorylation to determine if the improvement in learning is correlated with increased p-CREB. Consistent with earlier observations, we found that *APP^NL-G-F^*KI mice that received only saline injections had less p-CREB in the dentate gyrus than saline treated *APP^WT^* WT mice (p<0.01; Fig. 4, Supplemental Table 3). With PBA treatment, *APP^NL-G-F^* KI mice had increased phosphorylation of CREB compared to saline treated *APP^NL-G-F^* KI mice (p<0.05; Fig. 4, Supplemental Table 3). We next examined activation of AKT and quantified p-AKT using western blots and found that p-AKT levels were unchanged regardless of genotype or treatment (Fig. 4C, 4D). We found more hippocampal GADD34 in these late-stage *APP^NL-G-F^*KI saline-treated mice when compared to *APP^WT^* WT saline-treated mice (p<0.05; Fig. 4C’, 4F). PBA treatment led to a reduction in GADD34 in the *APP^NL-G-F^* KI mice, compared to *APP^NL-G-F^* KI saline-treated mice (p<0.001; Fig. 4C’, 4F). To determine if AMPA levels were changed with a late-stage treatment paradigm, we also probed for GluR1 via western blot. However, consistent with our other experimental paradigms, we observed no changes in GluR1 amount across groups. Interestingly, *APP^NL-G-F^* KI saline-treated mice did not display less hippocampal BDNF as compared to *APP^WT^* WT mice (Fig. 4C’’, 4G), yet late-stage PBA treatment resulted in increased BDNF in the *APP^NL-G-F^*KI mice (p<0.05; Fig. 4C’’, 4G). Collectively, these data suggest that even a late chaperone intervention can improve cognition, with corresponding increases in p-CREB and BDNF.

### 4: Late-stage PBA treatment reduces Aβ42 in *APP^NL-G-F^* KI mice

One of the key pathological features of AD is amyloid-β plaques that consist of insoluble Aβ fibrils (Chen et al., 2017; Sengupta et al., 2016). To determine if PBA treatment improved pathology, we carried out immunostaining using the 6E10 antibody, which labels Aβ plaques.

We did not observe any gross differences in plaque number in the hippocampi of early PBA and saline treatment *APP^NL-G-F^* KI mice (Fig. 5A), nor did we observe changes in plaque number in mice that received late-stage treatment (Fig. 5B). However, there is evidence that Aβ oligomers, as opposed to plaques, are the more toxic Aβ species (Chen et al., 2017; Sengupta et al., 2016). We probed Aβ42 levels using ELISAs in the late-stage older *APP^NL-G-F^* KI mice and found that PBA reduced the concentration of Aβ42 in the hippocampus of these mice compared to saline-treated *APP^NL-G-F^*KI mice (Fig. 5C). These results are consistent with the changes we observed in ADAM10 levels with PBA treatment, suggesting that modulating the UPR via PBA treatment can alter APP processing via increasing ADAM10.

**Fig. 5:**
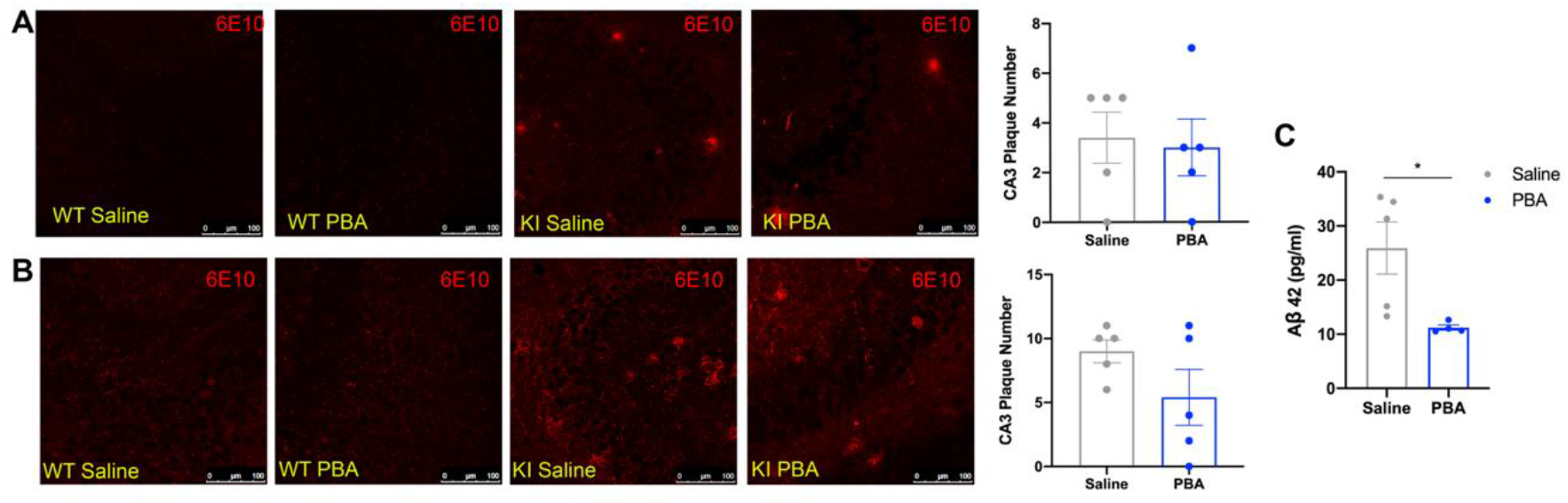
Late-stage PBA treatment in 10–12-month-old *APP^NL-G-F^* KI mice reduces Aβ-42 immunofluorescent images in the CA3 in the hippocampus. Confocal images of the hippocampus across groups. **A**: 6E10 immunofluorescent images in the CA3 in *APP^NL-G-F^* KI mice that received treatment from weaning; **B**: 6E10 immunofluorescent images in the CA3 in *APP^NL-G-F^*KI mice that received late-stage treatment; **C**: ELISA data using hippocampal homogenate from late-stage *APP^NL-G-F^* KI mice. (Data presented as mean ± SE; n=4-5 animals per group; T-test, *p<0.05).

### 5: Binding Immunoglobulin Protein (BiP) overexpression in the hippocampus of *APP^NL-G-F^* KI mice improves proteostasis and cognition

Having observed the effect of systemic chaperone administration on learning and UPR molecular markers we wanted to determine whether restoring chaperone levels in the hippocampus specifically would be sufficient to restore cognitive function and proteostasis in the *APP^NL-G-F^* KI mice. Using bilateral hippocampal stereotaxic surgical microinjections of an AAV-CaMKII-BiP we overexpressed the endogenous chaperone, BiP in the hippocampus of adult (6mos old) *APP^NL-G-F^* KI and WT littermate mice. An AAV-CaMKII-mCherry vector was used for control microinjections.

BiP overexpression was confirmed with immunofluorescence staining and western blotting (Fig. 6A, Supplemental Fig. 2, Supplemental Table 2). BiP was markedly increased in the CA3 of *APP^NL-G-F^* KI mice (p<0.001; Fig. 6A) and in the dentate gyrus of *APP^NL-G-F^* KI mice (p<0.01; Supplemental Figure 2, Supplemental Table 2). Control injections were also confirmed by mCherry staining in the CA3 and dentate gyrus (Fig. 6A, Supplemental Fig. 2).

**Fig. 6:**
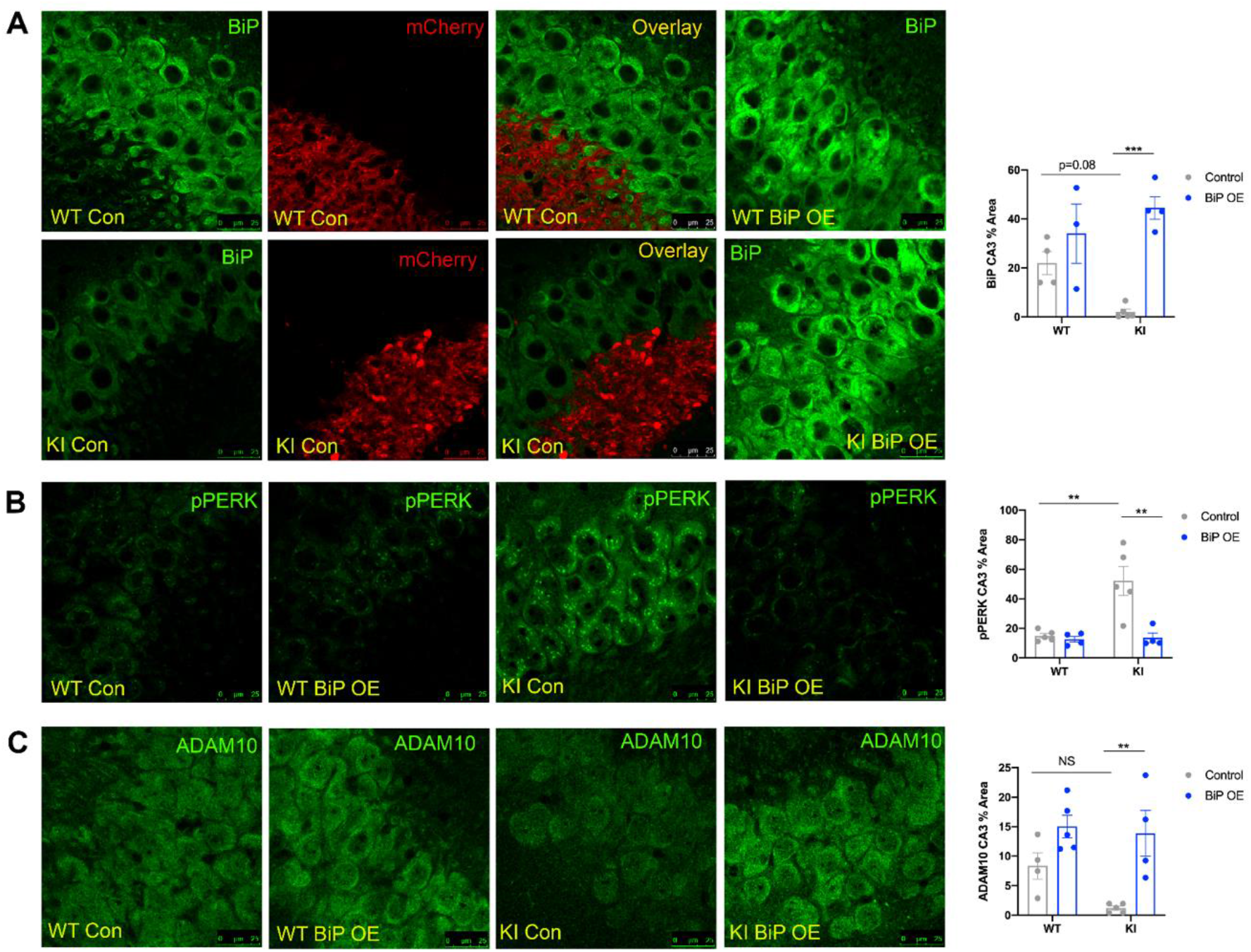
Local hippocampal AAV-BiP microinjections reduce PERK signaling and increase ADAM10 levels in *APP^NL-G-F^* KI mice. Confocal images of the hippocampal CA3 across groups. **A**: BiP staining and merged images shown for mCherry-tagged (red) control virus-injected mice; **B**: p-PERK; **C**: ADAM10; Mean ± SE percent area within hippocampal sections (n=3-5 animals per group; two-way ANOVA with Tukey post-hoc correction for multiple comparisons, *p<0.05, **p<0.01, ***p<0.001).

#### 5.1 Hippocampal BiP overexpression reduces PERK phosphorylation and increases ADAM10 in the APP^NL-G-F^ KI mice

Following confirmation of viral targeting and BiP overexpression, we probed for markers of UPR activation. *APP^NL-G-F^*KI mice that received control injections displayed more p-PERK staining in the CA3 than control-injected *APP^WT^* WT mice (p<0.01; Fig. 6B, Supplemental Table 2). With BiP overexpression, *APP^NL-G-F^* KI mice exhibited reduced p-PERK levels (p<0.01; Fig. 6B, Supplemental Table 2), indicating that overexpressing BiP directly into the hippocampus is sufficient to reduce PERK phosphorylation.

To determine if BiP overexpression was sufficient to promote non-amyloidogenic APP processing, we probed for ADAM10. With BiP overexpression, *APP^NL-G-F^* KI mice showed increased ADAM10 compared to control injected *APP^NL-G-F^* KI mice (p<0.01, Fig. 6C, Supplemental Table 2).

#### 5.2 Overexpressing BiP in the hippocampus of APP^NL-G-F^ KI mice improves learning

Next, we wanted to confirm if hippocampal BiP overexpression was sufficient to improve learning. We tested learning in *APP^NL-G-F^* KI and *APP^WT^* WT mice following BiP and control injections using the SOR and Y-maze tests. We found that the *APP^WT^*WT mice, regardless of injection, were able to successfully distinguish between the moved and unmoved objects in the SOR test (Fig. 7A, Supplemental Table 4). However, *APP^NL-G-F^* KI control injected mice were unable to differentiate between the objects compared to control injected *APP^WT^* WT mice (p<0.001; Fig. 7A, Supplemental Table 4). *APP^NL-G-F^*KI that received hippocampal BiP overexpression injections showed improved learning in the SOR test and were able to discriminate between the objects, compared to control injected *APP^NL-G-F^* KI mice (p<0.001; Fig. 7A, Supplemental Table 4). Again, there were no differences in how males and females performed in the test (Supplemental Fig. 4C-D). There were also no differences across groups in percent alternations in the Y-maze test (Fig. 7B). Overall, directly supplementing BiP levels in the hippocampus is sufficient to improve learning in the *APP^NL-G-F^* KI mouse model of AD.

**Fig. 7:**
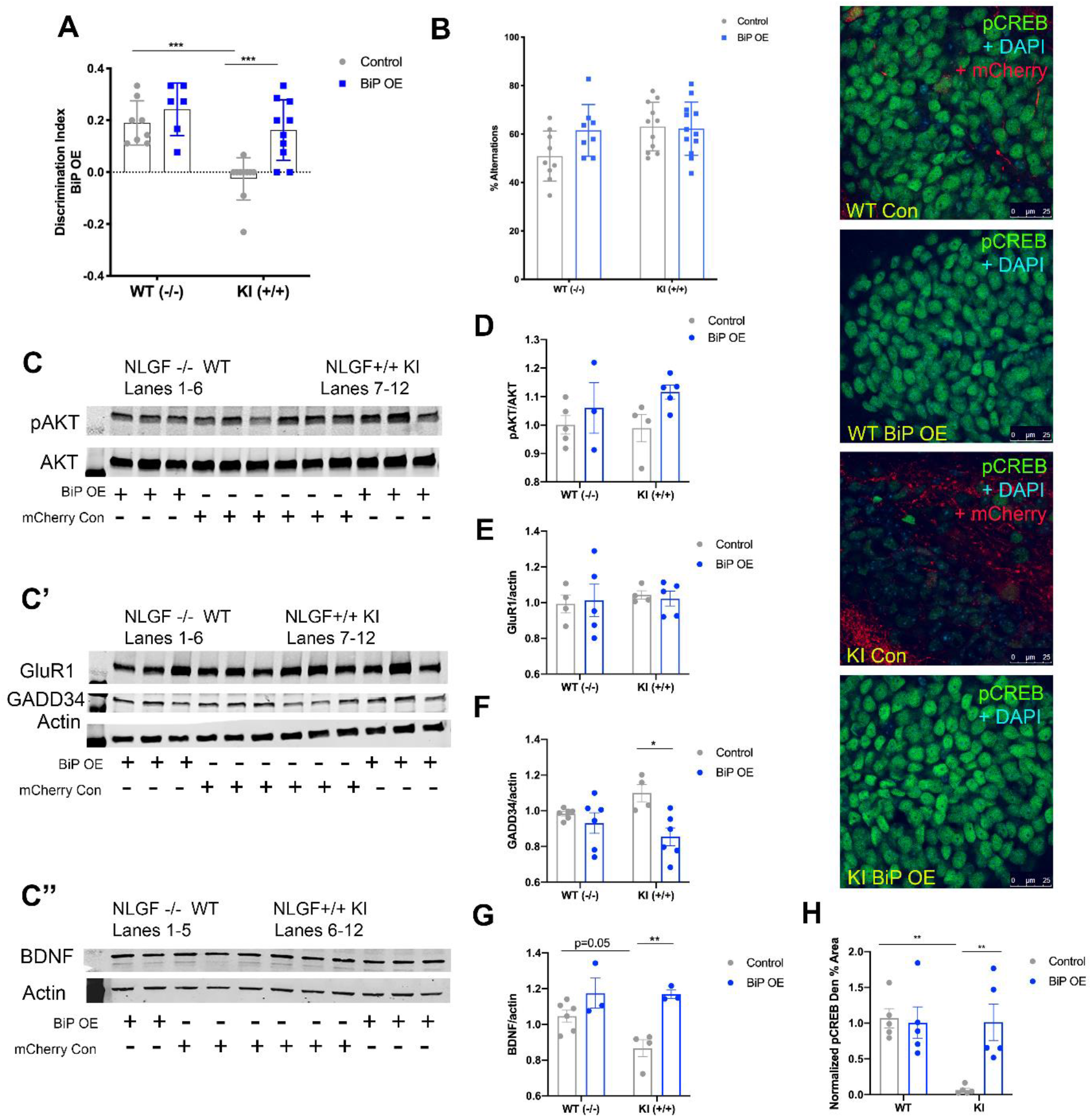
Local hippocampal AAV-BiP microinjection improves learning in the SOR test in *APP^NL-G-F^* KI mice, increases CREB phosphorylation, and improves BDNF levels. **A:** Discrimination index of SOR test; **B:** Percent alternations in the Y-maze test; **C-C’’:** Representative western blot images; each lane is a different animal. Membranes are cut along the MW markers following blocking to utilize the two-channel Odyssey system and varying molecular weights for the proteins of interest to probe for multiple markers within one membrane. **C:** pAKT and AKT, ∼65 kDa; **C’:** GluR1 ∼110 kDa, GADD34 ∼100 kDa, Actin ∼42 kDa; **C’’:** BDNF dimer ∼28 kDa, Actin ∼42 kDa. **D:** Quantification of western blot analysis for pAKT/AKT; **E:** Quantification of western blot analysis for GADD34/actin; **F:** Quantification of western blot analysis for GluR1/actin; **G:** Quantification of western blot analysis for BDNF/actin. **Right**: Confocal images of the hippocampal dentate gyrus across groups; **H**: Quantification of pCREB immunofluorescence normalized to *APP^WT^* WT mice, mean ± SE percent area within hippocampal dentate gyrus sections. n=3-6 animals per group for Western blot and n=5 for immunohistochemical assays. (All data presented as mean ± SE; all quantifications analyzed via two-way ANOVA with Tukey post-hoc correction for multiple comparisons, *p<0.05, **p<0.01, ***p<0.001).

#### 5.3 BiP overexpression induces CREB activation and BDNF levels in the APP^NL-G-F^ KI mice

We wanted to examine a mechanism for BiP-overexpression induced improvement in cognition. We probed for CREB phosphorylation in AAV-BiP and AAV-mCherry control-injected *APP^NL-G-F^* KI and *APP^WT^* WT mice. *APP^NL-G-F^* KI AAV-control injected mice displayed significantly less p-CREB in the dentate gyrus compared to *APP^WT^* WT controls (p<0.01; Fig. 7, Supplemental Table 2). With BiP overexpression injections, *APP^NL-G-F^* KI mice showed increased p-CREB compared to *APP^NL-G-F^*KI control injected mice (p<0.01; Fig. 7, Supplemental Table 2), suggesting that BiP overexpression in the hippocampus is sufficient to \ increase CREB phosphorylation. As in the other studies, we also examined p-AKT and GADD34 levels. We found that AKT phosphorylation did not significantly change across groups (Fig. 7C, 7D). In addition, *APP^NL-G-F^* KI control injected mice did not display an increase in GADD34 compared to control-injected *APP^WT^* WT littermates (Fig. 7C’, 7F). However, BiP overexpression in the *APP^NL-G-F^*KI mice led to a reduction in GADD34 levels, compared to control-injected *APP^NL-G-F^*KI mice (p<0.05; Fig. 7C’, 7F). Further, as with PBA and saline-treated mice, GluR1 levels did not change, regardless of genotype or injection (Fig. 7C’, 7E). However, *APP^NL-G-F^* KI control injected mice exhibited less BDNF levels as compared to *APP^WT^* WT control-injected mice (p=0.05; Fig. 7C’’, 7G) and BiP overexpression led to an increase in BDNF in *APP^NL-G-F^* KI mice (p<0.01; Fig. 7C’’, 7G). Altogether, these results demonstrate that local hippocampal BiP overexpression is sufficient to improve learning in these *APP^NL-G-F^* KI mice, through increasing CREB phosphorylation and BDNF.

## DISCUSSION

In this study, we postulated that utilizing chaperone treatment in an APP KI mouse model of AD would improve proteostasis and cognition. Encouragingly, our results indicate that chaperone therapy is sufficient to reduce chronic UPR activity and improve cognitive performance in the SOR test in the *APP^NL-G-F^* mouse model of AD. In particular, the control-treated *APP^NL-G-F^*KI mice displayed reduced levels of XBP1s, regardless of age, which is restored with PBA treatment. Importantly, changes in XBP1s levels are linked to APP processing, as XBP1s is known to promote a disintegrin and metalloproteinase 10 (ADAM10) (Peron et al., 2018; Reinhardt et al., 2014). ADAM10 is an alpha-secretase that cleaves APP in a non-amyloidogenic manner (Yuan et al., 2017). In AD, ADAM10 levels are reduced and APP is preferentially cleaved by BACE1, a beta-secretase, and gamma-secretases, ultimately generating Aβ42 in the amyloidogenic pathway (Sogorb-Esteve et al., 2018; Yuan et al., 2017). Under healthy conditions, APP has a functional role in the synapse, involved in cell adhesion and synapse stability (Hick et al., 2015; Muller et al., 2012). We show here that ADAM10 levels are reduced in *APP^NL-G-F^* KI mice, but are restored with PBA treatment, correlated with the changes we observed in XBP1s staining. Thus, modulating the UPR via chaperone therapy has a direct impact on APP processing.

We did not observe any changes in the number of plaques with PBA treatment, but this is not inconsistent with literature showing that Aβ plaques are not the most toxic Aβ species (Chen et al., 2017; Ondrejcak et al., 2010; Sengupta et al., 2016). Indeed, Aβ plaques have been observed in the brains of non-AD cognitively normal patients and animals, indicating that larger aggregates do not cause cognitive impairments and that Aβ oligomers are the more toxic species of Aβ (Erten-Lyons et al., 2009; Mormino and Papp, 2018; Rodrigue et al., 2009; Sloane et al., 1997). On the other hand, soluble Aβ oligomers are known to lead to cognitive deficits even in the absence of plaques (Gandy et al., 2010; Sengupta et al., 2016). While we saw no changes in plaques, we observed a reduction in Aβ42 with PBA treatment in late-stage intervention *APP^NL-G-F^* KI mice. These results are consistent with the changes observed in ADAM10, indicating that modulating the UPR can impact Aβ42 amounts. However, while ADAM10 levels were increased with chaperone treatment, ADAM10 itself has been associated with long-term depression (LTD) and synaptic downscaling (Gardoni et al., 2012; Malinverno et al., 2010). Therefore, the changes we observed in ADAM10 could be linked to improved pathology but do not fully explain the improvement in cognition with chaperone treatment. It is likely that a combination of factors, including the increase in p-CREB and BDNF together with reduced Aβ42 as a result of increased ADAM10 contribute to the improved learning phenotype.

Supplementing chaperone levels to modulate the UPR affects other cellular processes, including those involved in memory formation. Promisingly, the impaired cognition observed in the *APP^NL-G-F^*KI saline-treated and AAV-control-injected mice was rescued by PBA and BiP, respectively. Importantly, our data indicate that PBA treatment, even administered at a late stage in disease progression, was sufficient to restore cognition in the *APP^NL-G-F^* KI mice. Our findings align with a recently published study that demonstrates the crucial role of endoplasmic reticulum genes, especially that of BiP, in long-term memory (Chatterjee et al., 2022). Similar to our observations Chatterjee et al found that BiP overexpression rescued long-term memory deficits in a tau-based mouse model of ADRD. That study along with data presented here highlights the significance of chaperone proteins in the consolidation of memory. While in the present study we utilized the SOR test, future experiments should conduct a repertoire of tasks, including fear conditioning or the Morris Water Maze test in order to corroborate our findings.

We have previously shown that supplementing chaperone levels in aged mice reduces PERK activation, derepressing global protein translation which is necessary for memory formation. In this aging model, we showed that chaperone treatment reduced PERK signaling and GADD34 levels, leading to an increase in CREB activation, correlated with improved learning in the aged mice (Hafycz et al., 2022). GADD34 upregulation prevents phosphorylation of AKT, a CREB kinase, thus impairing CREB phosphorylation and thus, cognition (Farook et al., 2013; Sen, 2019). In the present study, we examined if a similar mechanism was involved in this mouse model of AD. Overall, our results show similar changes in GADD34 and p-AKT with PBA treatment and BiP overexpression, suggesting that a mechanism comparable to what we previously identified in a model of aging could apply to this *APP^NL-G-F^*KI model of AD, indicating that some age-related cellular processes may be involved in AD progression. Further, it is known that in AD there is a reduction of AMPA receptor levels that is linked to impaired cognition (Guntupalli et al., 2016; Martin-Belmonte et al., 2020). Here, we report no change in the GluR1 subunit of AMPA receptors via western blot using whole-hippocampal homogenate. However, this method lacks regional specificity, thus inhibiting the measure of potential local hippocampal differences in AMPA levels. Many of the immunohistochemical assays revealed hippocampal region-specific changes, and thus it is possible that GluR1 levels also change in hippocampal subregions. A future direction for this work would be to examine synaptosomal preparations, which could inform more nuanced changes in AMPA distribution directly at the synapse.

Under normal conditions, the UPR functions as an adaptive response to acute ER stress (Hetz et al., 2020; Ron and Walter, 2007). In the short term, PERK leads to attenuated translation, while ATF6 and IRE1 lead to increased chaperone synthesis and activation of protein degradation pathways (Hetz et al., 2020; Ron and Walter, 2007). However, in the presence of unresolved ER stress, chronic UPR activity initiates pro-apoptotic signaling through the PERK-CHOP pathway (Hetz et al., 2020; Lin et al., 2007). Indeed, it has been shown that the pro-survival arms of the UPR, namely IRE1-induced XBP1 splicing that promotes chaperone synthesis, is attenuated during persistent ER stress (Lin et al., 2007). This change is critical as it is thought to serve as a turning point for the cell from survival mechanisms to the initiation of apoptosis. In the case of neurodegenerative diseases, and AD specifically, proteostasis is chronically dysfunctional, likely contributing to the pathological aggregations observed and the progression of neurodegeneration (Halliday and Mallucci, 2014; Hetz and Saxena, 2017; Hoozemans et al., 2009; Hoozemans et al., 2012; Hughes and Mallucci, 2019). Interestingly, we observed reduced levels of BiP and XBP1s in the *APP^NL-G-F^* KI mice, which were rescued with PBA treatment from weaning and late-stage intervention. This suggests that supplementing chaperone levels, even later in disease progression, promotes pro-survival UPR activation, as enhancing XBP1s levels could increase protein folding capacity by promoting endogenous BiP. Further, late-stage intervention saline-treated *APP^NL-G-F^* KI mice displayed an increase in CHOP compared to age-matched *APP^WT^* WT saline-treated mice, suggesting that if left untreated, this APP KI mouse model of AD leads to UPR-related pro-apoptotic signaling. CHOP levels were reduced with PBA treatment, again providing evidence that supplementing chaperone levels promotes an adaptive UPR response and reduces chronic UPR signaling.

ER stress is believed to play an important role in AD. Other animal models of AD display ER stress, however because these models overexpress membrane proteins, like APP or PS1, ER stress may be induced non-specifically as these proteins can be misfolded and aggregate in the ER (Hashimoto et al., 2018). Thus, the use of the *APP^NL-G-F^* KI model, which increases Aβ without overexpressing APP, provides a unique tool to discern if ER stress in these AD models is due to Aβ pathology as opposed to overexpressed APP. Interestingly, the research group that developed the *APP^NL-G-F^* KI model for AD also examined ER stress in these mice (Hashimoto et al., 2018). The authors claim that there is no induction of ER stress in these *APP^NL-G-F^* KI mice and thus that ER stress observed in other AD models is nonspecific (Hashimoto et al., 2018).

However, those observations are limited by low animal numbers and Western blot assays using whole brain region homogenates, overlooking potential effects in subregions or specific neural populations. There is a robust volume of literature that has examined ER stress in mouse models and post-mortem human tissue and has reported increased ER stress, including increased PERK activation (Hetz and Saxena, 2017; Hoozemans et al., 2005; Ma et al., 2013; Unterberger et al., 2006). It is therefore vitally important to distinguish effects that are due to specific disease pathology, as opposed to changes resulting from the limitations inherent in the animal models used (i.e. APP overexpression). We observed increases in several markers of ER stress in the *APP^NL-G-F^* KI saline-treated and control-injected mice, which could be due in part by utilizing higher animal numbers as well as immunofluorescence to examine hippocampal subregions.

It is also important to note that PBA, in addition to acting as a protein chaperone, also functions as a histone deacetylase (HDAC) inhibitor (Cuadrado-Tejedor et al., 2013; Ricobaraza et al., 2009). Our data which focuses on the chaperone function of PBA has been substantiated by Chatterjee et al (2022) who showed that sodium butyrate which acts solely as an HDAC inhibitor does not rescue cognition in an AD model. Further substantiating a role for chaperones our data also indicates that local hippocampal BiP overexpression was sufficient to improve both memory and proteostasis in *APP^NL-G-F^*KI mice. These experiments corroborate the role of PBA as a chaperone on proteostasis and cognition and exemplify the role of ER stress in behavior.

Altogether, the results presented here are consistent with an extensive body of literature that has studied the role of UPR signaling in pathology and memory in AD (Halliday and Mallucci, 2014; Hetz and Saxena, 2017). PERK signaling is known to be increased in tissue of AD patients as well as in various animal models of AD (Halliday and Mallucci, 2014; Hetz and Saxena, 2017). Critically, several studies have shown that reducing UPR activity or modulating the UPR improves memory in AD models (Halliday et al., 2015; Moreno et al., 2012; Ricobaraza et al., 2012). It has been shown that inhibiting phosphorylation of eIF2a, downstream of PERK and responsible for attenuating global translation, improves memory and synaptic plasticity in models of AD (Halliday et al., 2017; Moreno et al., 2012; Oliveira et al., 2021). Another study demonstrated that ATF4 is linked to impaired memory and synaptic plasticity (Costa-Mattioli et al., 2009). In addition, other studies have linked XBP1s to BDNF (Cisse et al., 2017; Martinez et al., 2016). BDNF is a growth factor known to promote memory formation (Bekinschtein et al., 2008; Gonzalez et al., 2019; Panja and Bramham, 2014), and its expression is impaired in AD, which is thought to be linked to the hallmark cognitive impairments (Amidfar et al., 2020; Ng et al., 2019; Tanila, 2017).

Thus, we propose that there are multiple pathways that are beneficially modulated by chaperone treatment as is illustrated in Figure 8. Supplementing chaperone levels, by increasing XBP1s, serves to ameliorate disease-related characteristics in several ways. The first is that XBP1s drives ADAM10 to reduce aberrant Aβ42. XBP1s also leads to improved memory-formation via promoting BDNF, as well as improved proteostasis by promoting synthesis of the chaperone BiP. Meanwhile, PERK signaling is reduced with both PBA treatment and local hippocampal BiP overexpression, suggesting that the protein translation block via peIF2α is lifted, allowing for the normal translation to proceed. Phosphorylated CREB levels are also increased with chaperone treatment, which could in part be due to a reduction in PERK-ATF4- GADD34 signaling. In the late-stage intervention paradigm, PERK-CHOP signaling is ameliorated, suggesting that UPR-related pro-apoptotic signaling is reduced. It is these collective changes that shift the cell towards adaptive pro-survival signaling, making chaperone supplementation an appealing target for the treatment of AD.

**Figure 8:**
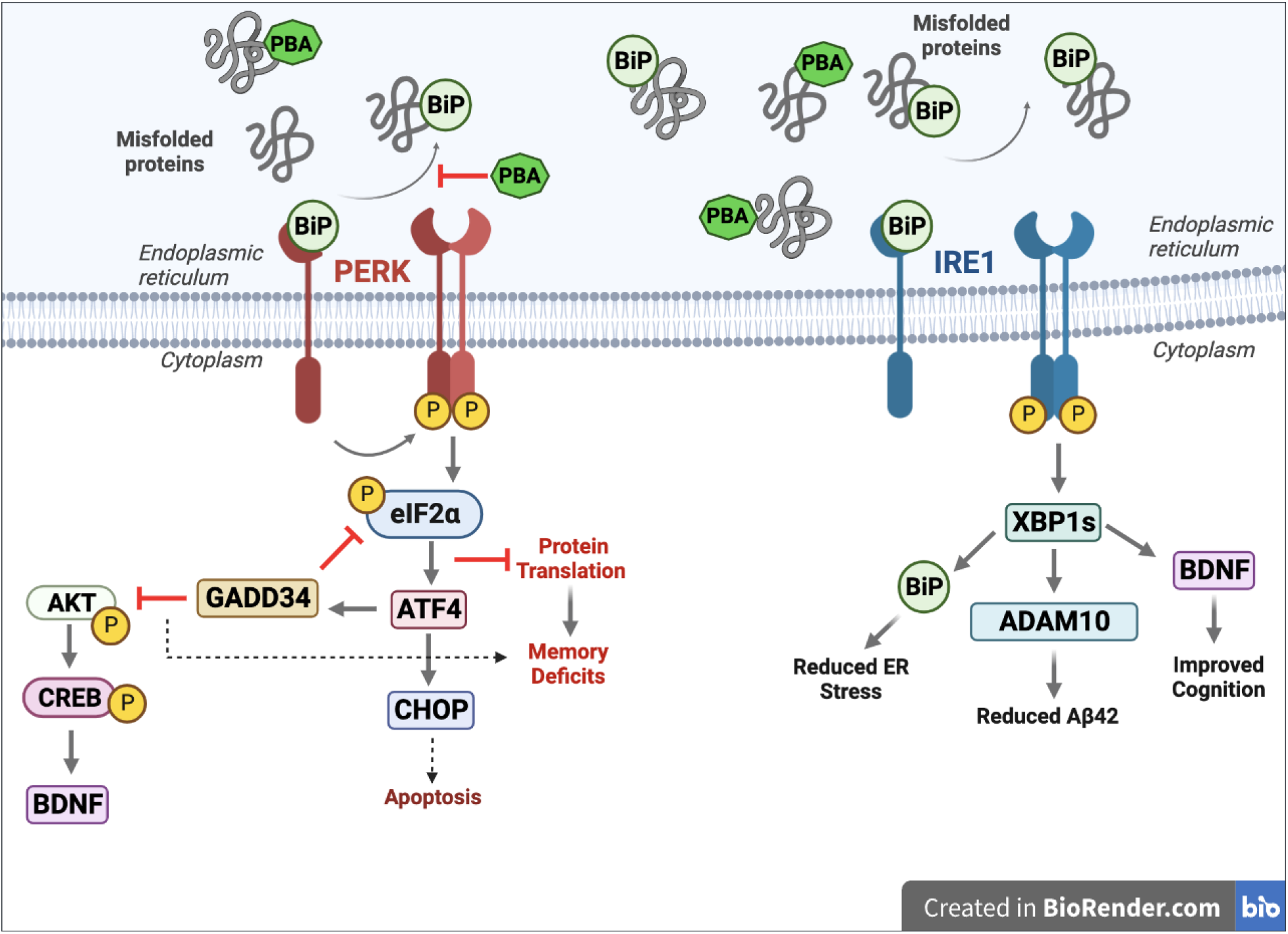
Summary figure. Perk activation following BiP dissociation leads to inhibited protein translation via p-eIF2a, increased GADD34, reduced p-CREB signaling and increased CHOP. Chaperone therapy reduces PERK activation and promotes IRE1 activation, which leads to increased XBP1s, BiP, ADAM10, and BDNF.

## CONCLUSION

Our results show that supplementing protein chaperone levels improves proteostasis and cognition in this *APP^NL-G-F^*KI mouse model of Alzheimer’s disease. Chaperone treatment leads to an adaptive UPR response that is linked to non-amyloidogenic APP processing and increased CREB phosphorylation. Encouragingly, even late-stage chaperone treatment improves cognition and proteostasis. Collectively, this work could provide valuable insight into the development of novel therapeutics for this debilitating and pervasive disease.

## MATERIALS AND METHODS

### Mice

Animal experiments were conducted in accordance with the guidelines of the University of Pennsylvania Institutional Animal Care and Use Committee. Mice were maintained as previously described (Chellappa et al., 2019; Naidoo et al., 2008). Mice were housed at 23°C on a 12:12 h light/dark cycle and had ad libitum access to food and water. *APP^NL-G-F^* KI and *APP^WT^* WT mice were used for these experiments (Saito et al., 2014).

For PBA treatment studies two groups of mice were used and a third group of mice were used for AAV-BiP overexpression and AAV-mCherry control injections. Animal numbers for each genotype, sex, and treatment condition used for SOR cognitive testing can be found in Supplemental Table 5.

### Drug administration

Sodium 4-phenylbutyrate (PBA) (Cayman Chemical, Ann Arbor, MI) was administered as previously described (Hafycz et al., 2022). Briefly, PBA was given in twice weekly via intraperitoneal (IP) injections at a dose of 40mg/kg and in the drinking water as a 0.8% PBA concentration in a 1% sucrose solution. Vehicle-treated mice received IP sterile saline injections twice weekly and were given a 1% sucrose solution as drinking water. Treatment began when mice were 4-6 weeks of age (from weaning) or 10-12 months of age (late intervention) and continued for 10-12 weeks. Following treatment, all underwent cognitive behavioral testing.

Mice were then sacrificed via transcardial saline perfusions at ZT0 immediately following the recovery sleep period. Half brains were collected and either flash frozen or fixed in 4% paraformaldehyde.

### Stereotaxic Injection Surgery

Stereotaxic hippocampal injections were performed to deliver a BiP viral overexpression vector (Vector Biolabs; AAV5-CamKIIa-GRP78; titer 4.8×10^12^ GC/ml) and an mCherry control vector (Addgene; pAAV5-CaMKIIa-mCherry; titer 2.3 x10^-13^ GC/ml) into the hippocampi of young and aged mice as previously described (Hafycz et al., 2022). Following general anesthesia (2% isoflurane) and sterilization of surgical area and tools, an incision was made on the top of the skull and local bupivacaine was administered. Four burr holes were drilled for bilateral hippocampal injections, with coordinates as follows: AP ±1.8mm, ML ± 0.8mm, ±1.8mm, DV - 1.7mm, -1.9mm. Using a 1 µL Hamilton Syringe, 50 nanoliters of virus was injected per burr hole. Mice were given subcutaneous injections of meloxicam and saline post-operation for analgesia and hydration, respectively. Topical antibiotic ointment was used following the suturing of the surgical site. Mice were observed for 3 days of recovery. Behavioral testing occurred four weeks after surgery date to allow for sufficient viral expression. Transcardial perfusions were performed at ZT0 following testing, and half brains were collected and either flash frozen or fixed in 4% paraformaldehyde.

### Spatial Objection Recognition (SOR) Test

The Spatial Objection Recognition test is well-established hippocampal-dependent spatial memory test (Bevins and Besheer, 2006; Cavoy and Delacour, 1993; Hafycz et al., 2022). The mice are placed together for an hour in the testing apparatus for three consecutive days prior to testing to acclimate to the container. Testing occurs in two phases. The first is the training phase where two identical objects are placed on one side of the apparatus. Mice are placed individually in the apparatus for 10 minutes and all interactions (smelling, touching, etc.) are counted. After training, the mice are returned to their home cage and left undisturbed for about one hour. Before the start of the testing phase, one of the identical objects is moved to the opposite side of the apparatus. Mice are placed in the apparatus during the testing phase similar to the training phase, but only for 3 minutes. Interactions with each object are measured again. Discrimination index calculations were performed as previously described as a measure for how well the mice distinguish between the moved and unmoved object (Sivakumaran et al., 2018).

### Y-Maze Spontaneous Alternation Test

The Y-Maze test was performed as previously described (Kraeuter et al., 2019). Briefly, a single mouse was placed in the center of the apparatus and was allowed to move freely through the maze for 5 minutes. Each individual arm entry and the order in which the entries occurred were recorded. After testing, the number of alternations (3 separate arm sequential arm entries) were counted and presented as a percentage.

### Immunohistochemical Assays

Post-fixed half-brain coronal sections were sliced at 40 μm using a cryostat as previously described (Hafycz et al., 2022; Zhu et al., 2007). Every other section was placed in 24-well plates containing cryoprotectant for free floating immunohistochemistry staining and stored at - 20°C, as previously described (Naidoo et al., 2008; Naidoo et al., 2018). For all markers, we compared n=4-8 in each of the four groups.

### Immunofluorescence (IF)

Immunofluorescence staining was conducted as previously described, with a n=4-8 animals per group per assay (Hafycz et al., 2022; Naidoo et al., 2011). Primary antibodies are as follows: pCREB (ser133) (1:300, Cell Signaling 87G3); CREB (1:200, Cell Signaling 86B10); pPERK (Thr980) (1:200, Bioss bs-3330R); ATF4 (1:500, ProteinTech, 60035-1-Ig); peIF2α (1:100, Cell Signaling 3597S); BiP/anti-KDEL (1:1000, Enzo Life Sciences ADI-SPA-827F). Secondary antibodies are as follows: Alexa Fluor 488 donkey anti-rabbit IgG (1:500); Alex Fluor 594 donkey anti-mouse IgG (1:500); Alexa Fluor 488 donkey anti-mouse IgG (1:500).

### Quantitative Analysis of IF Images

Confocal images were acquired as previously described (Hafycz et al., 2022; Owen et al., 2021), using Leica SP5/AOBS microscope. Confocal laser intensities, nm range, detector gain, exposure time, amplifier offset, and depth of the focal plane within sections per antigen target were standardized across compared sections. Confocal images were quantified as previously described (Zhu et al., 2018). Briefly, 3-4 sections were imaged per animal (n=4-5 animals per group).

Using ImageJ software, the images were converted to an 8-bit grayscale with detection threshold standardized across all images to detect percent areas. The percentage area covered within the target region was measured and average percent areas for each mouse were analyzed.

### Western Blot Staining

Frozen brain tissue was prepared for western blot assays as previously described (Hafycz et al., 2022; Naidoo et al., 2008; Naidoo et al., 2018). Briefly, brain tissue was homogenized on ice with lysis buffer containing protease inhibitors. After centrifugation, protein concentration for each sample was determined with a BCA protein assay and samples were prepared such that each contained 20µg of protein. SDS-PAGE gels were run as previously described (Naidoo et al., 2008), and protein bands were imaged and quantified via infrared imaging on an Odyssey scanner (LiCor). For all markers we compared n=3-8 for each of the groups. Primary antibodies are as follows: GADD34 (1:500, Protein Tech 10449-1-AP); pAKT (1:500, Cell Signaling 9271); Akt (1:500, Cell Signaling 9272); BDNF (1:500, Abcam ab108319); β-Actin (1:2000, Santa Cruz sc-47778); GluR1 (1:500, Novus NBP2-61472). Secondary antibodies are as follows: LiCor IRDye 680RD Goat anti-Mouse (1:10,000); LiCor IRDye 800RD Goat anti-Mouse (1:10,000); LiCor IRDye 800RD Goat anti-Rabbit (1:10,000); Odyssey IRDye 680 Goat anti-Rabbit (1:10,000).

### Statistical Analyses

Data are presented as the average ± standard error of the mean (SEM) of sample size n. Statistical analyses were performed in PRISM (GraphPad Software, La Jolla, CA). Two-way ANOVA was used to determine interaction effects, with Tukey post-hoc corrections for multiple comparisons. p<0.05 was the threshold for determining statistical significance. In some instances, as interaction tests are typically underpowered, regardless of the significance of the ANOVA term, we subsequently performed specific between group comparisons defined by *APP^NL-G-F^* genotype and treatment. We evaluated whether outcome measures differed between genotype and treatment using two-tailed T-tests (see supplement). Together, this set of analyses provides a more comprehensive evaluation of the *APP^NL-G-F^* genotype and treatment group relationships.

## List of Supplementary Materials

**Figure S1:**
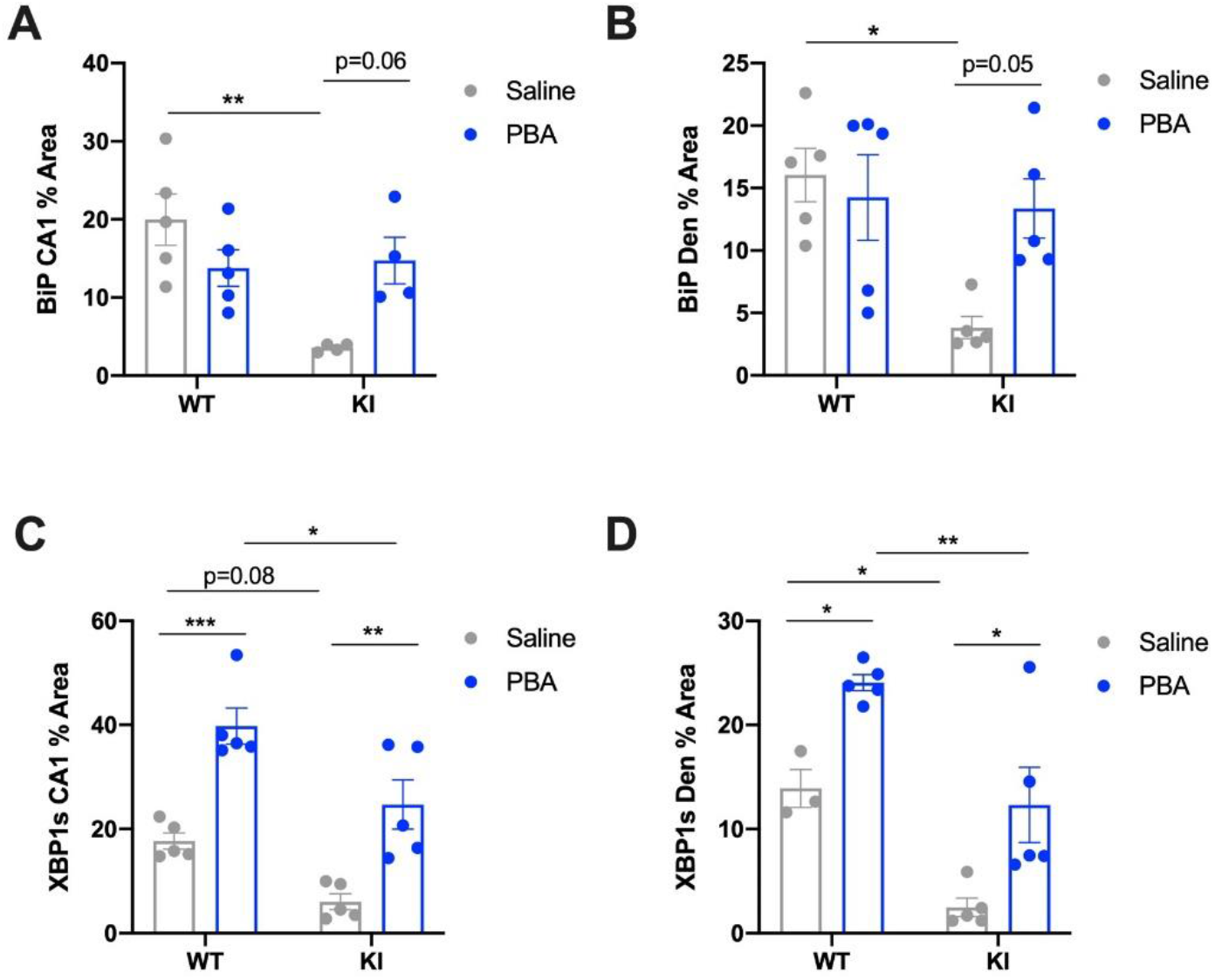
PBA treatment from weaning increases BiP and XBP1s in the hippocampus of *APP^NL-G-F^* KI mice. Immunofluorescence quantification of BiP and XBP1s in hippocampal subregions. A: BiP in CA1; B: BiP in dentate gyrus; C: XBP1s in the CA1; D: XBP1s in dentate gyrus. (n=3-5 animals per group; two-way ANOVA with Tukey post-hoc correction for multiple comparisons, *p<0.05, **p<0.01, ***p<0.001).

**Figure S2.**
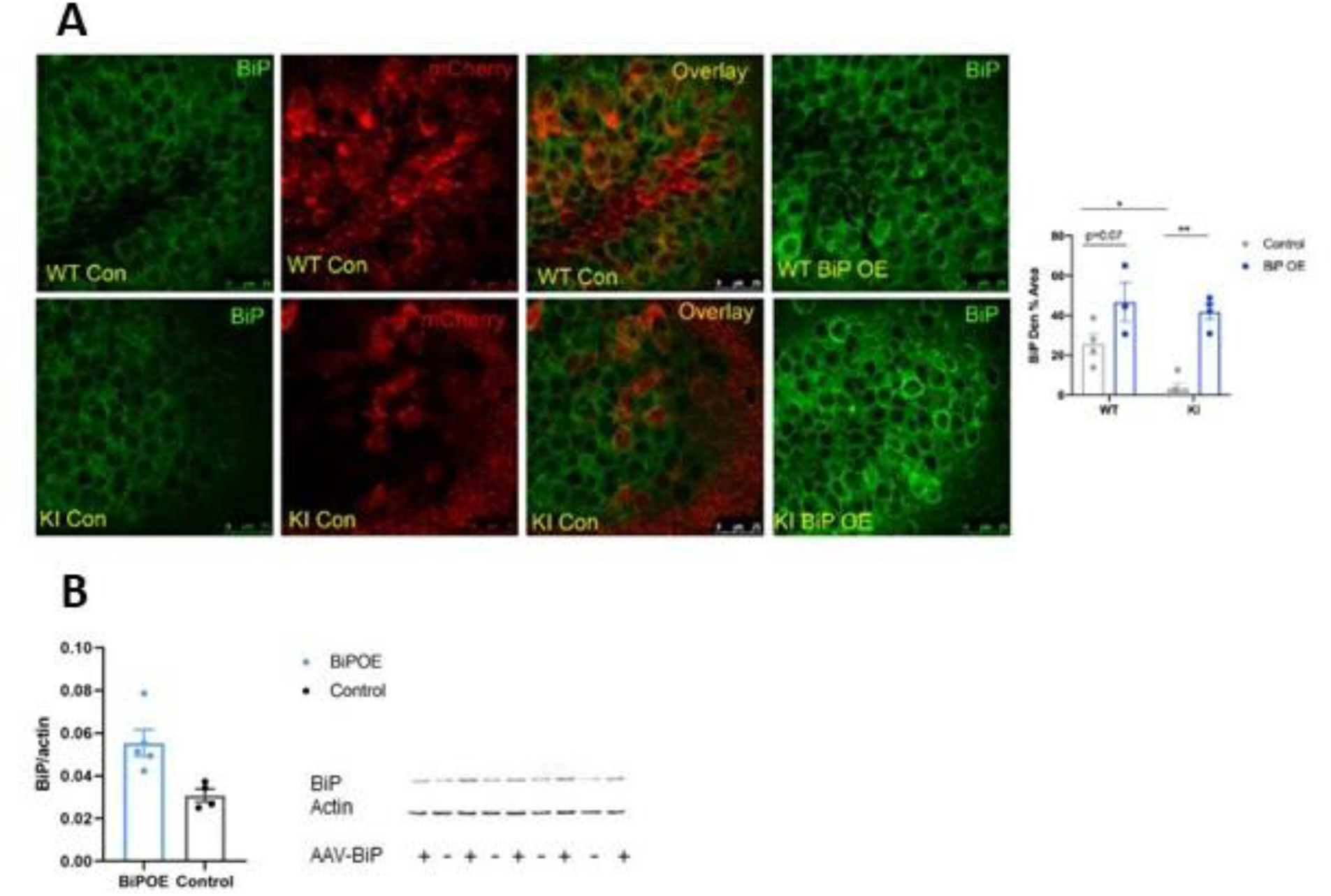
A: Quantification of BiP immunofluorescence in the dentate gyrus following local hippocampal AAV-BiP and AAV-mCherry microinjections. Representative confocal images of the dentate gyrus across groups. Top Row: *APP^WT^* WT mice Bottom Row: *APP^NL-G-F^* KI mice; Right: Quantification of immunofluorescence in the dentate gyrus. Mean ± SE percent area of BiP within hippocampal dentate gyrus sections (n=4-5 animals per group; two-way ANOVA with Tukey post-hoc correction for multiple comparisons, *p<0.05, **p<0.01). **B:** western blot quantification of BiP in KI bulk hippocampus and representative image following AAV-BiP microinjections.

**Figure S3:**
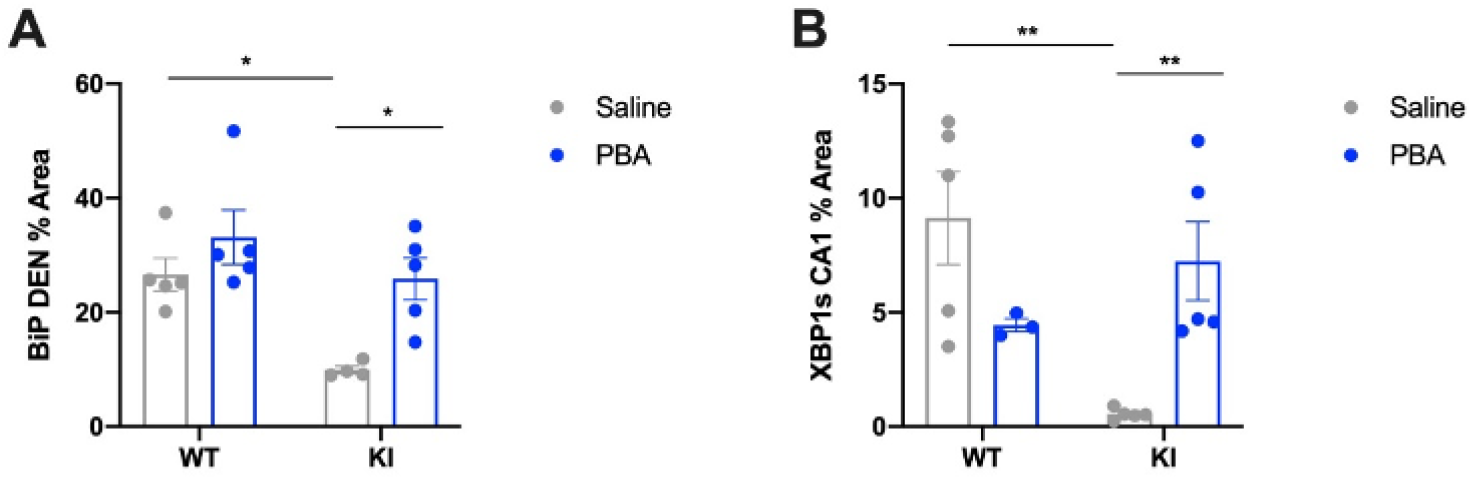
Late-stage PBA treatment increases BiP and XBP1s in 10–12-month-old *APP^NL- G-F^* KI mice. Immunofluorescence quantification of BiP and XBP1s in hippocampal subregions. A: BiP in dentate gyrus; B: XBP1s in CA1. (n=3-5 animals per group; two-way ANOVA with Tukey post-hoc correction for multiple comparisons, *p<0.05, **p<0.01).

**Figure S4:**
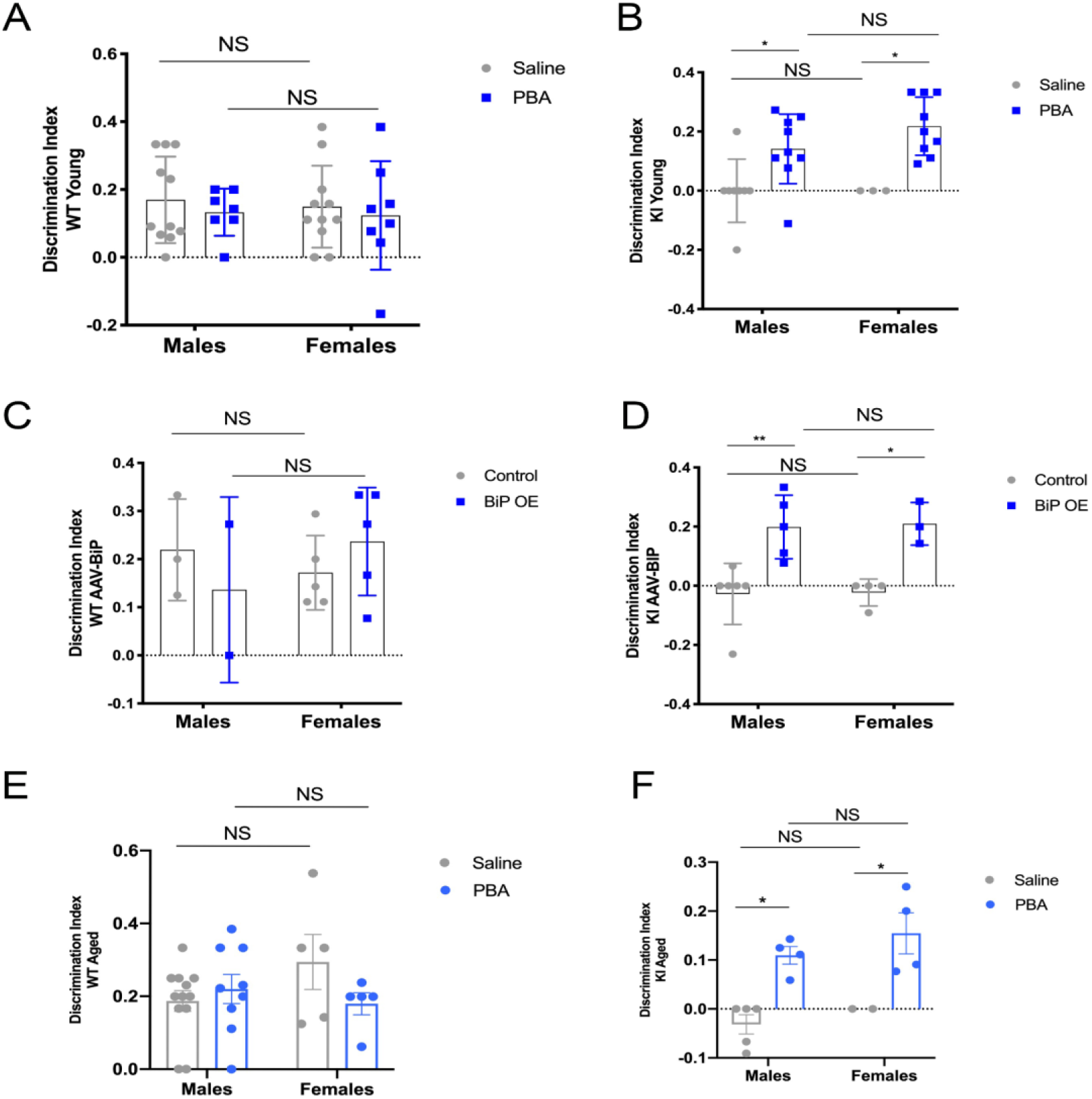
SOR discrimination index did not vary by sex in any chaperone treatment paradigm. Discrimination index of SOR test by sex across groups. A-B: Discrimination index in mice following PBA treatment from weaning; A: *APP^WT^* WT mice; B: *APP^NL-G-F^* KI mice; C-D: Discrimination index in mice following AAV-BiP or AAV-mCherry hippocampal microinjections; C: *APP^WT^* WT mice; D: *APP^NL-G-F^* KI mice; E-F: Discrimination index in mice following late-stage PBA treatment; E: *APP^WT^*WT mice; F: *APP^NL-G-F^* KI mice. (All analyses performed using two-way ANOVA with Tukey post-hoc correction for multiple comparisons, *p<0.05, **p<0.01).

**Table S1:** ANOVA interaction values for immunofluorescence assays in *APP^NL-G-F^* KI and *APP^WT^* WT mice following saline and PBA treatment from weaning.

| **<u>Marker</u>** | **<u>P<sub>int</sub></u>** |
| --- | --- |
| BiP | **CA1: 0.005** |
|  | **CA3: 0.0009** |
|  | **DEN: 0.03** |
| pPERK | CA1: 0.61 |
|  | CA3: 0.61 |
|  | DEN: 0.69 |
| ATF4 | **CA1: 0.0002** |
|  | CA3: 0.87 |
|  | **DEN: 0.008** |
| XBP1 | CA1: 0.59 |
|  | CA3: 0.88 |
|  | DEN: 0.95 |
| ADAM10 | **CA1: 0.0003** |
|  | **CA3: 0.0001** |
|  | **DEN: 0.003** |
| p-CREB | **DEN: 0.05** |
| **<u>Significant values (p<0.05) in bold</u>** |  |

**Table S2:** ANOVA interaction values for immunofluorescence assays in *APP^NL-G-F^* KI and *APP^WT^* WT mice following AAV-mCherry and AAV-BiP injections.

| Marker | p<sub>int</sub> |
| --- | --- |
| BiP | **CA3: 0.02** |
|  | DEN: 0.12 |
| pPERK | **CA3: 0.008** |
|  | **DEN: 0.02** |
| ADAM10 | CA3: 0.20 |
|  | DEN: 0.31 |
| p-CREB | **DEN: 0.01** |
| Significant values (p<0.05) in bold |  |

**Table S3:** ANOVA interaction values for immunofluorescence assays in 10–12-month-old mice *APP^NL-G-F^* KI and *APP^WT^* WT mice following late-stage saline and PBA intervention.

| Marker | p<sub>int</sub> |
| --- | --- |
| BiP | **CA1: 0.03** |
|  | **CA3: 0.03** |
|  | DEN: 0.21 |
| pPERK | CA1: 0.97 |
|  | CA3: 0.70 |
|  | DEN: 0.68 |
| ATF4 | CA1: 0.35 |
|  | CA3: 0.87 |
|  | DEN: 0.52 |
| CHOP | CA1: 0.97 |
|  | **CA3: 0.05** |
|  | DEN: 0.62 |
| XBP1 | **CA1: 0.002** |
|  | **CA3: <0.0001** |
|  | **DEN: 0.01** |
| ADAM10 | CA1: 0.57 |
|  | **CA3: 0.03** |
|  | DEN: 0.59 |
| p-CREB | *DEN: 0.08* |
| Significant values ($p < 0.05$) in bold | |

**Table S4:**
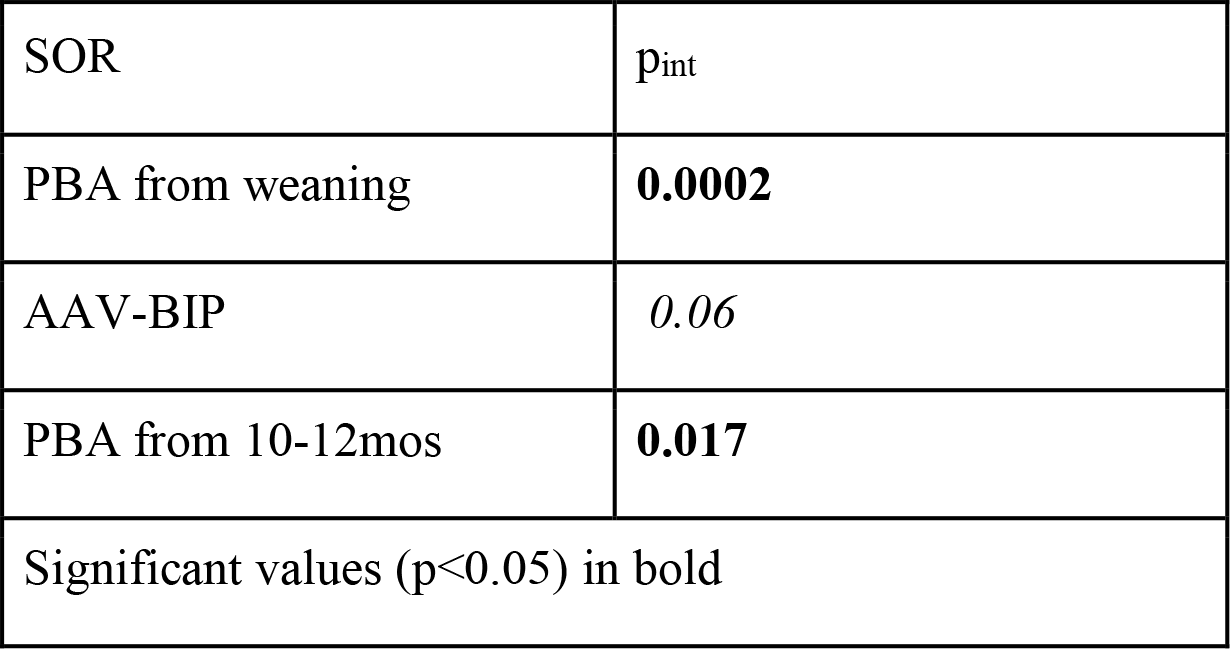
ANOVA interaction values for the discrimination index across SOR tests.

**Table S5:** Mouse number distributions across all SOR experiments.

|  | Treatment from Weaning |  |  |  |
| --- | --- | --- | --- | --- |
|  | Saline |  | PBA |  |
|  | Males | Females | Males | Females |
| WT | 11 | 11 | 7 | 8 |
| KI | 7 | 3 | 9 | 9 |
|  | AAV-BiP and AAV-mCherry |  |  |  |
|  | AAV-mCherry |  | AAV-BiP |  |
|  | Males | Females | Males | Females |
| WT | 3 | 5 | 1 | 5 |
| KI | 6 | 4 | 5 | 5 |
|  | Late-Stage Treatment |  |  |  |
|  | Saline |  | PBA |  |
|  | Males | Females | Males | Females |
| WT | 12 | 5 | 9 | 6 |
| KI | 6 | 2 | 4 | 6 |
| Table of total mouse numbers utilized across SOR behavioral experiments. |  |  |  |  |

## Acknowledgments

The authors wish to acknowledge and thank Mr. Robert Komlo for assistance with weekly I.P. injections, and to Mr. Brendan Keenan of the Sleep Statistical Core for assistance with statistical analyses.

## Author contributions

JH designed and conducted the experiments, analyzed data, and wrote the manuscript. ES conducted western blot experiments. NN conceptualized and designed the study, provided the resources, interpreted data, wrote and edited the manuscript.

## Data and materials availability

The raw data supporting the conclusions in this manuscript will be made available upon request.

